# Intestinal lipid metabolism controls immune response through NHR-68 and gut–brain signaling in *C. elegans*

**DOI:** 10.64898/2026.06.24.734305

**Authors:** Jie Ren, Yu Sang, Young-Mo Kim, Ernesto S. Nakayasu, Alejandro Aballay

## Abstract

Animals must allocate limited energetic resources across competing defense programs in response to infection. Here, we show that the conserved nuclear hormone receptor NHR-68 integrates fatty acid metabolism with the neural control of molecular and behavioral immunity in *Caenorhabditis elegans*. Acting in parallel with NHR-10, NHR-68 controls genes involved in polyunsaturated fatty acid (PUFA) metabolism. Loss of NHR-68 disrupts linoleic acid (LA) homeostasis, impairing pathogen avoidance behavior. Supplementation with LA restores avoidance, and *fat-3* inhibition, which elevates LA, enhances pathogen avoidance, whereas loss of LA synthesis by *fat-2* inhibition diminishes this behavior, indicating that LA promotes behavioral immunity. We further show that NHR-68 acts in the intestine to regulate linoleic acid homeostasis, and that changes in intestinal lipid metabolism influence an AWC-dependent pathogen-avoidance circuit through intestine-to-neuron communication. NHR-68 suppresses activation of the PMK-1/p38 MAPK and DAF-16/FOXO pathways, which mediate molecular immune responses. These findings identify a gut-brain transcriptional circuit that connects intestinal lipid metabolism to neural and immune outputs, revealing a mechanism by which the metabolic state coordinates behavioral and molecular defenses to optimize host protection.

## Introduction

Animals must continuously balance the allocation of limited resources among competing physiological needs, including growth, metabolism, and defense. Mounting immune responses is energetically costly, and excessive or prolonged activation can compromise fitness. As a result, host defense systems have evolved strategies that integrate immune signaling with metabolic state and environmental cues to optimize protection while preserving energy homeostasis. The coordination of these regulatory layers across tissues remains poorly understood.

Nuclear hormone receptors (NHRs) are a family of transcription factors that couple metabolic state to gene-expression programs controlling development, metabolism, reproduction, and stress responses [1]. These receptors act by binding to specific signaling molecules, such as steroid hormones, retinoids, and thyroid hormones, which modulate gene expression to control a variety of physiological processes [2]. In *C. elegans*, NHRs are involved in key biological functions such as responses to oxidative stress [3], starvation [4, 5], hypoxia [6], and pathogen infection [7–12]. The *C. elegans* genome encodes over 280 NHRs, more than in humans or mice, and only a small subset has been functionally characterized [13] .

In *C. elegans*, the intestine is a major metabolic and immune organ that also influences behavior through inter-tissue communication [14–17]. In addition to molecular immune responses, *C. elegans* relies on pathogen avoidance as an important behavioral defense strategy. Animals can detect pathogenic bacteria and undergo aversive learning to reduce subsequent exposure, and this process is regulated by multiple neuronal and signaling pathways [18–20]. Fatty-acid metabolism is a central readout of intestinal metabolic state governed by networks of NHRs [21, 22]. *C. elegans* has evolved an array of conserved desaturase (*fat*) and elongase (*elo*) genes that can synthesize a broad range of fatty acids de novo from acetyl-CoA [23]. In addition to their roles in nervous system function [24], both monounsaturated fatty acids (MUFAs) and polyunsaturated fatty acids (PUFAs) can exhibit antibacterial functions [25]. The MUFA oleate is necessary for innate immune gene expression and protects *C. elegans* from *Pseudomonas aeruginosa* infection [26, 27]. The PUFA eicosapentaenoic acid (EPA) also enhances the *C. elegans* immune response and functions as an effector that can disrupt or inhibit biofilm formation, helping to regulate host immunity against *Candida albicans* [28]. Genetic studies have shown that fatty-acid desaturation and elongation pathways, including *fat-2* and *fat-3*, regulate polyunsaturated fatty acid composition and affect susceptibility to *P. aeruginosa* infection [27, 29]. In addition, fatty-acid supplementation can modulate host immune responses and survival during bacterial infection, supporting a functional link between lipid metabolism and innate immunity [26, 30]. Despite these links, the specific intestinal metabolic changes that occur during bacterial infection and how they are integrated with neural control remain unclear.

Here, we show that NHR-68, a member of the expanded *C. elegans* nuclear hormone receptor family with similarities to mammalian hepatocyte nuclear factor 4-alpha (HNF4α), acts in parallel with NHR-10 to coordinate fatty acid metabolism, innate immune signaling, and pathogen avoidance. NHR-68 regulates genes involved in PUFA biosynthesis, and its loss reduces linoleic acid (LA), a fatty acid that promotes avoidance behavior. We found that NHR-68 acts in the intestine and signals through the AWC chemosensory neuron to drive behavioral immunity, while suppressing the PMK-1/p38 MAPK and DAF-16/FOXO pathways that mediate molecular immune responses.

Together, these findings identify a transcriptional mechanism that links intestinal metabolism to neural and immune outputs. Finally, our results suggest a general principle for cross-tissue coordination of defense. By aligning intestinal lipid metabolism with neural avoidance and modulation of immune activation, nuclear receptors provide a way to balance energetic costs and benefits during infection. This principle may be conserved given the broad conservation of nuclear receptor signaling and lipid-derived cues.

## Results

### NHR-10 and NHR-68 inhibit innate immune activation

Nuclear hormone receptors (NHRs) are well-characterized regulators of metabolism, development, and reproduction[1], but their involvement in innate immune regulation remains poorly understood. To identify NHRs that control immune activation, we performed a reverse genetic screen using 236 *nhr* genes from the Ahringer *C. elegans* RNAi feeding library [31, 32]. As a readout of intestinal immune activation, we used a transcriptional reporter for *clec-60,* a C-type lectin gene whose expression is strongly induced in response to infection and other immune triggers [33–35]. The *clec-60p::gfp* reporter is widely used as a readout for the activation of protective innate immune pathways in the *C. elegans* intestine [33–35].

We identified nine *nhr* genes (*nhr-77*, *nhr-10*, *nhr-130*, *nhr-222, nhr-162, nhr-42, nhr-140*, *nhr-68*, and *nhr-54*) whose knockdown resulted in pronounced increased *clec-60p::gfp* expression (Fig 1A, Fig S1 and Table S1). Most of these NHRs, including *nhr-77*, *nhr-10*, *nhr-42*, *nhr-68*, and *nhr-54*, have been identified in genome-wide transcription factor profiling studies as components of regulatory networks associated with metabolic and stress-responsive processes [36], although their roles in intestinal immune regulation remain largely uncharacterized. Notably, *nhr-10* and *nhr-68* were of particular interest, as they are known to function together as a persistence detection module in propionate metabolism [37]. Given their shared roles in metabolic regulation and in the expression of immune gene *clec-60*, we focused our analysis on *nhr-10* and *nhr-68*. Animals treated with double RNAi targeting both *nhr-10* and *nhr-68* showed a further enhancement of the *clec-60p::gfp* signal compared to either single knockdown (Fig 1A and Fig S1). This was confirmed by qRT-PCR, which revealed an ∼8–10-fold increase in *clec-60* transcript levels in single knockdowns and a ∼20-fold increase in the double knockdown (Fig 1B), indicating that the two NHRs have additive effects in suppressing immune activation.

**Fig 1.**
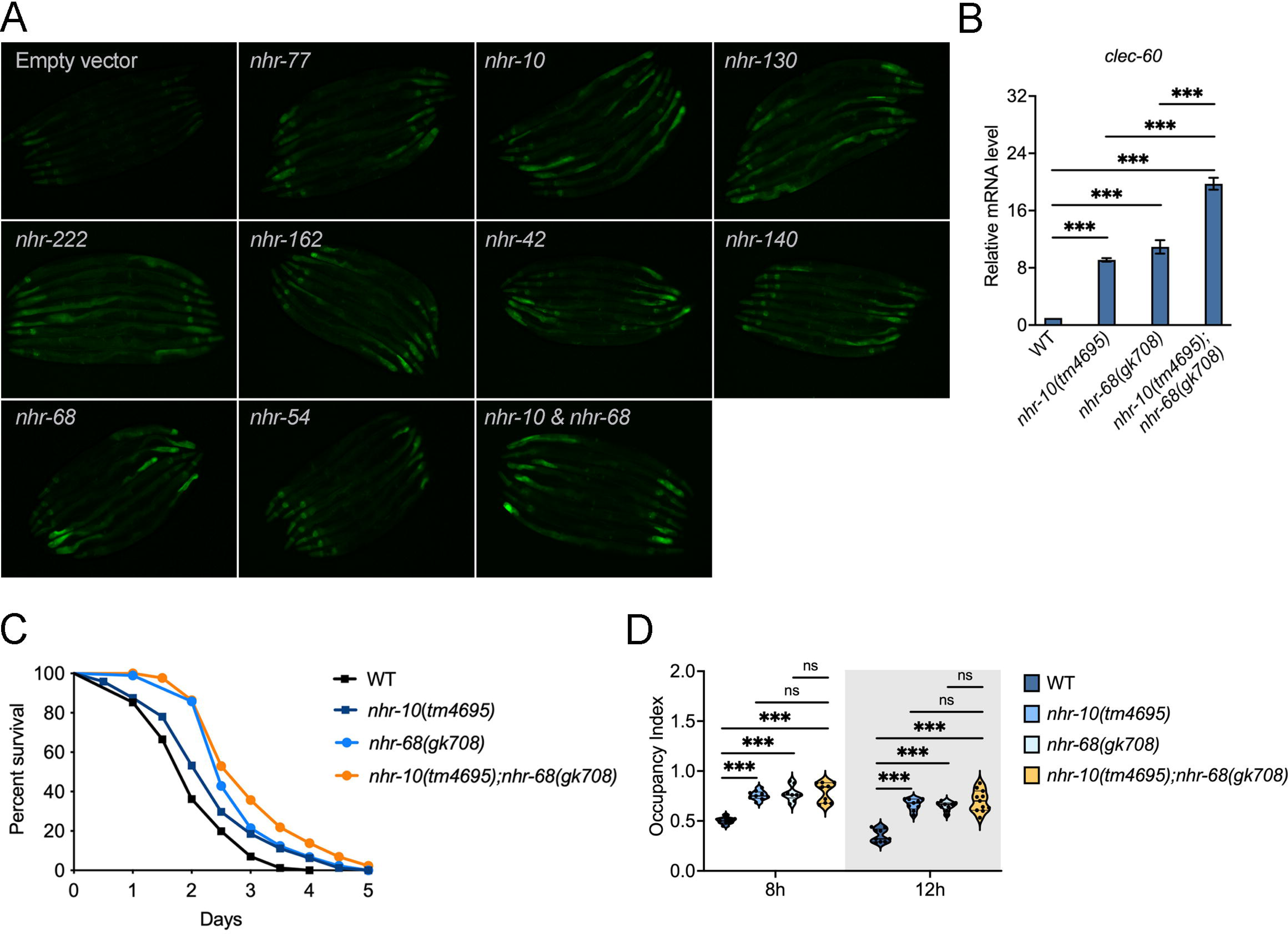
NHR-10 and NHR-68 inhibit innate immune response. (A) Representative fluorescence micrographs of *clec-60p::gfp* animals grown on *E. coli* HT115 carrying empty vector or RNAi against *nhr-77*, *nhr-10*, *nhr-130*, *nhr-222*, *nhr-162*, *nhr-42*, *nhr-140*, *nhr-68*, or *nhr-10* and *nhr-68* (n = 8 animals per condition per experiment, three independent experiments). Fluorescence images were captured with 100-ms exposures. Scale bar, 200 µm. (B) qRTÖPCR analysis of the immune gene *clec-60* in wild-type, *nhr-10(tm4695)*, *nhr-68(gk708)* and *nhr-10(tm4695)*; *nhr-68(gk708)* animals fed on *E. coli*.

Data are presented as the means ± SDs from three independent experiments, n = 200 per condition. \*\*\**p* < 0.001, one-way ANOVA with Tukey’s multiple-comparison test. (C) Survival of WT, *nhr-10(tm4695)*, *nhr-68(gk708)* or *nhr-10(tm4695)*;*nhr-68(gk708)* animals grown on *P. aeruginosa* at 25 °C. Mean survival times were 2.06, 2.36, 2.82, and 3.13 days, respectively. Data are presented as means from three independent experiments, n = 90 per condition. WT versus *nhr-10(tm4695)*, *p* < 0.01; WT versus *nhr-68(gk708)*, *p* < 0.001; WT versus *nhr-10(tm4695)*; *nhr-68(gk708)*, *p* < 0.001; *nhr-10(tm4695)* versus *nhr-10(tm4695)*; *nhr-68(gk708)*, *p* < 0.001; *nhr-68(gk708)* versus *nhr-10(tm4695)*; *nhr-68(gk708)*, *p* =0.0435, the Kaplan-Meier method was used to calculate the survival fractions, and statistical significance between survival curves was determined using the log-rank test. (D) Occupancy index for WT, *nhr-10(tm4695)*, *nhr-68(gk708)* or *nhr-10(tm4695)*;*nhr-68(gk708)* animals on *P. aeruginosa* at 8 hr and 12 hr. The occupancy index was calculated as (N _on_ lawn/N _total_).

We then evaluated the *nhr-10(tm4695)*, *nhr-68(gk708)*, two deletion alleles predicted to cause loss of function, as well as *nhr-10(tm4695);nhr-68(gk708)* animals for susceptibility to *P. aeruginosa*. Both single mutants survived longer than wild-type animals, indicating increased resistance, and *nhr-10(tm4695);nhr-68(gk708)* animals survived slightly longer than either single mutant, consistent with an additive effect of the two genes (Fig 1C). *C. elegans* uses behavioral avoidance as a defense against pathogens [38, 39]. We found that *nhr-10(tm4695)* and *nhr-68(gk708)* animals showed significantly reduced bacterial avoidance behavior, with a modest further reduction in the *nhr-10(tm4695);nhr-68(gk708)* animals (Fig 1D and S2A). Locomotion assays confirmed that the reduced avoidance phenotype was not due to impaired locomotor ability in these mutants (Fig S2B). These findings indicate that NHR-10 and NHR-68 concurrently suppress molecular immunity linked to pathogen resistance while promoting pathogen avoidance behavior.

Because *nhr-68(gk708)* animals displayed a pathogen-resistant phenotype, reduced bacterial avoidance, and increased *clec-60* expression, we next asked whether the expression of *nhr-68* could reverse these phenotypes. To address this, we constructed an *nhr-68p::nhr-68::gfp* translational reporter expressing GFP-tagged NHR-68 under its native promoter to rescue expression in the mutant background. As shown in Fig 2A, *nhr-68::gfp* was expressed in the hypodermis, intestine, and head neurons, consistent with a previous study [40]. Expression of *nhr-68* also rescued the increased *clec-60* mRNA levels in *nhr-68(gk708)* animals (Fig 2B) and partially rescued pathogen susceptibility (Fig 2C) and bacterial avoidance (Figs 2D and S2C). These results indicate that *nhr-68* expression compensates for the mutant phenotype and that *nhr-10* and *nhr-68* act in a parallel fashion to control molecular and behavioral immune responses to *P. aeruginosa*.

**Fig 2.**
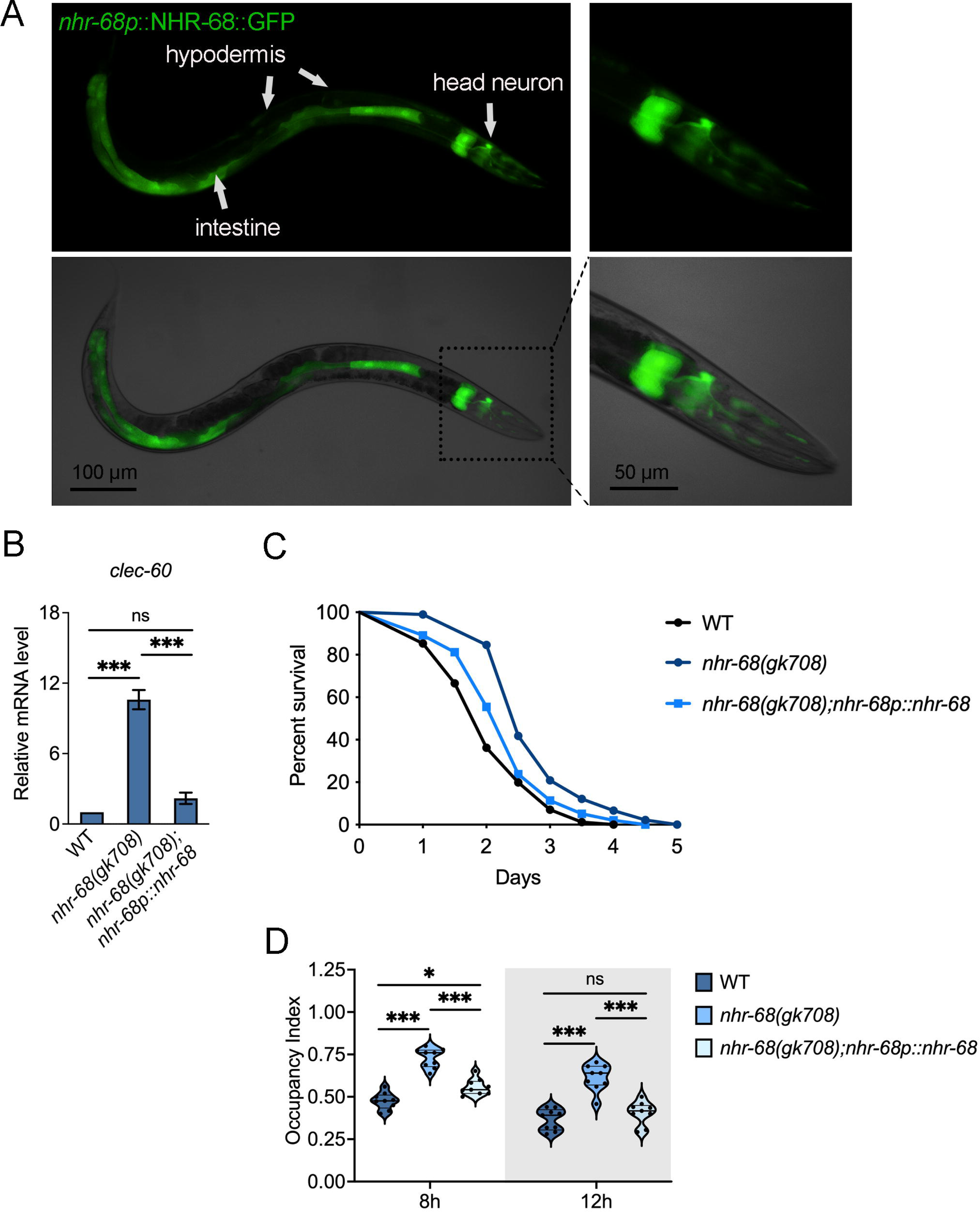
The expression of *nhr-68* restored pathogen susceptibility. (A) Representative fluorescence micrographs of *nhr-68(gk708);nhr-68p::nhr-68::gfp* animals. n = 30; data are representative of four independent experiments. Fluorescence images were captured using 400-ms exposures. Scale bars indicate 100 µm in the left two panels and 50 µm in the right two panels. (B) qRTÖPCR analysis of the immune gene *clec-60* in wild-type, *nhr-68(gk708)* or *nhr-68(gk708);nhr-68p::nhr-68* animals fed on *E. coli*. Data are presented as the means ± SDs from three independent experiments, n = 200 per condition. \*\*\**p* < 0.001, one-way ANOVA with Tukey’s multiple-comparison test. (C) Survival of wild-type N2, *nhr-68(gk708)* or *nhr-68(gk708);nhr-68p::nhr-68* animals on *P. aeruginosa* at 25 °C. Mean survival times were 2.07, 2.83, and 2.35 days, respectively. Data are presented as means from three independent experiments, n = 90 per condition. Wild-type versus *nhr-68(gk708)*, *p* < 0.0001; Wild-type versus *nhr-68(gk708); nhr-68p::nhr-68*, *p* = 0.0002; *nhr-68(gk708)* versus *nhr-68(gk708); nhr-68p::nhr-68*, *p* =0.0202. (D) *P. aeruginosa* occupancy indexes for WT, *nhr-68(gk708)* and *nhr-68(gk708);nhr-68p::nhr-68* animals at 8 hr and 12 hr after seeding. The occupancy index was calculated as (N _on_ lawn/N _total_).

Previous studies showed that NHR-10 and NHR-68 regulate genes linked to vitamin B12 metabolism and act together as a persistence detector for propionate metabolism, enabling metabolic network rewiring in response to dietary inputs [37]. To assess whether vitamin B12 supplementation affects the NHR-68–regulated immune response, we measured expression of the immune gene *clec-60* with or without vitamin B12. Vitamin B12 did not alter *clec-60* expression in *nhr-68(gk708)* animals (Fig S3A), suggesting that the NHR-68-mediated regulation of *clec-60* is unlikely to be directly linked to vitamin B12 metabolism under these conditions.

We showed that *nhr-68(gk708)* animals exhibit increased pathogen resistance against *P. aeruginosa* (Figs 1C and 2C). To test whether vitamin B12 influences pathogen resistance, we examined resistance using *E. faecalis*, which unlike *P.* aeruginosa, does not synthesize vitamin B12 [41] [42]. As shown in Fig S3B, *nhr-68(gk708)* animals displayed increased resistance to *E. faecalis*, and this phenotype was unaffected by vitamin B12 supplementation. Together with the additive effects observed for NHR-10 and NHR-68 in controlling *clec-60* expression and pathogen resistance (Figs 1C and 1D), these results support a model in which NHR-10 and NHR-68 both contribute to the regulation of molecular and behavioral immune responses.

### NHR-68 inhibits the PMK-1 and DAF-16 pathways during immune activation

To investigate the mechanism underlying the increased survival of *nhr-68*(*gk708*) animals upon *P. aeruginosa* (Fig 1C), we performed an RNA-seq analysis of the changes in gene expression profiles of these animals after 12 h infection with *P. aeruginosa*. Principal component analysis (PCA) of the transcriptomes revealed clear separation between *nhr-68(gk708)* and wild-type animals, indicating distinct gene expression patterns (Fig S4A). To reduce false positives, only genes with a fold change greater than two and *p* < 0.05 were considered differentially expressed (Table S2).

An unbiased gene enrichment analysis performed using the Database for Annotation, Visualization, and Integrated Discovery (DAVID) (https://david.ncifcrf.gov/)[43] revealed that the innate immune response Gene Ontology (GO) cluster had the second-highest number of hits, following signal transduction. Other enriched GO clusters corresponded to transmembrane transport and proteolysis (Fig 3A and Table S3).

**Fig 3.**
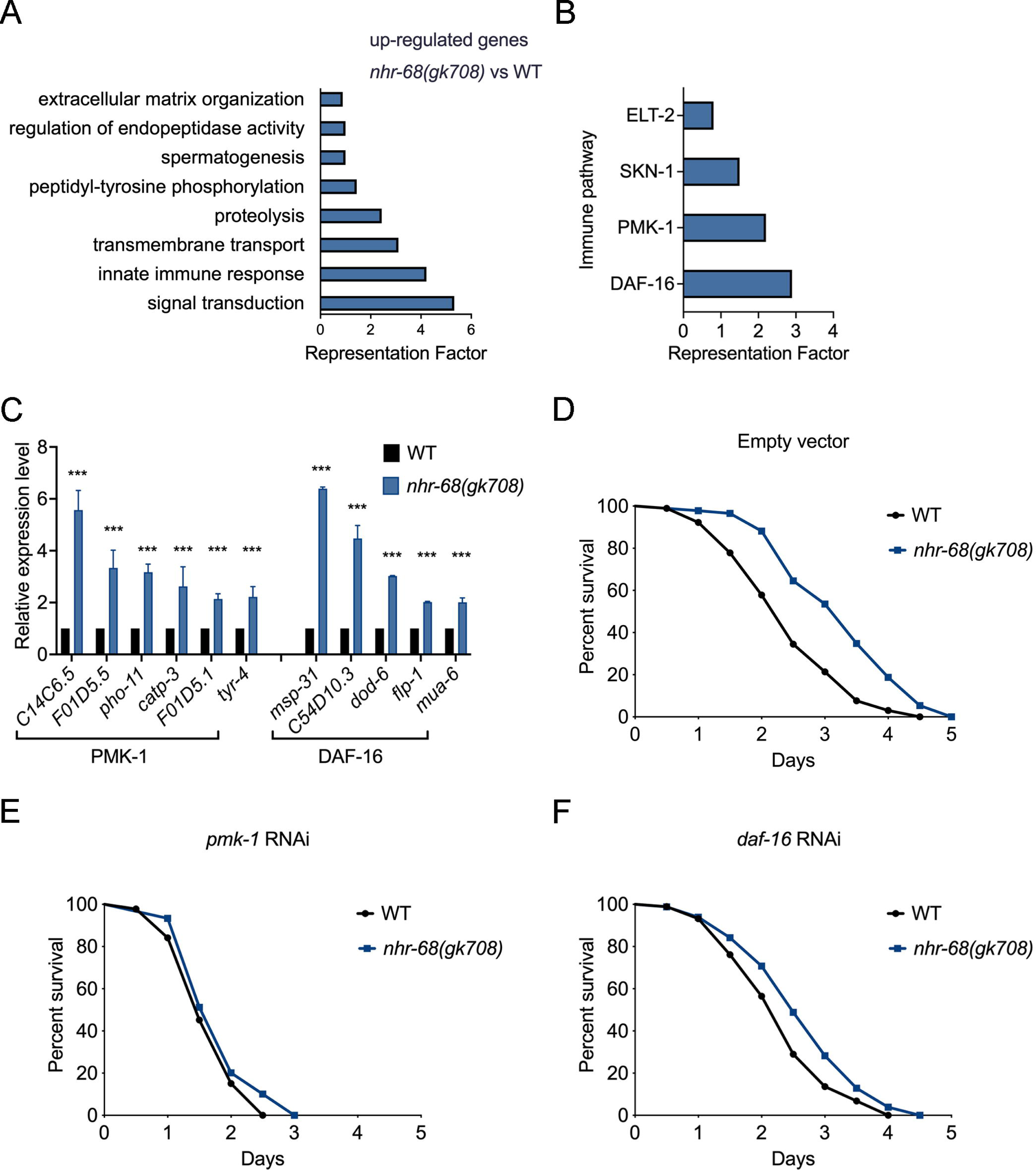
NHR-68 inhibits immune pathways. (A) Gene Ontology analysis of genes upregulated in *nhr-68(gk708)* versus WT animals after 12h infection with *P. aeruginosa*. The results were filtered to include only terms significantly enriched in *nhr-68(gk708)* with a q value <0.1. (B) The representation factors of the identified upregulated genes controlled by the PMK-1, DAF-16, and SKN-1 pathways. (C) qRTOPCR analysis of PMK-1 and DAF-16-dependent genes in *nhr-68(gk708)* compared with WT animals (n = 3; three technical replicates in each case). (D) Survival of wild-type and *nhr-68(gk708)* animals fed empty vector control RNAi followed by exposure to *P. aeruginosa*. Mean survival times were 2.40 and 3.01 days, respectively. Wild-type versus *nhr-68(gk708)*, *p* < 0.0001. (E) Survival of wild-type and *nhr-68(gk708)* animals fed *pmk-1* RNAi followed by exposure to *P. aeruginosa*. Mean survival times were 1.88 and 1.92 days, respectively. Wild-type versus *nhr-68(gk708)*, *p* = 0.1121. (F) Survival of wild-type and *nhr-68(gk708)* animals fed *daf-16* RNAi followed by exposure to *P. aeruginosa*. Mean survival times were 2.14 and 2.43 days, respectively. Wild-type versus *nhr-68(gk708)*, *p* = 0.0130. Panels D–F are shown separately to enable direct comparison of survival under control, *pmk*-1 RNAi, and *daf-16* RNAi conditions; statistical comparisons were performed between wild-type and *nhr-68(gk708)* animals within each RNAi condition, and no direct statistical comparisons between RNAi treatments were performed.

To further explore immune activation, we compared genes upregulated in *nhr-68(gk708)* animals to previously identified gene sets regulated by immune pathways in *C. elegans* (Table S4). Gene sets controlled by PMK-1 and DAF-16 showed significant overlap with genes whose expression increased in *nhr-68(gk708)* animals infected with *P. aeruginosa* (Fig 3B). qRT–PCR validation of selected genes confirmed the RNA-seq data (Fig 3C), supporting the conclusion that loss of *nhr-68* leads to activation of these immune genes.

To identify NHR-68-controlled genes in the absence of infection, we also performed RNA-seq on *nhr-68(gk708)* and wild-type animals fed only on *E. coli* OP50. PCA analysis showed clear separation between the two groups, indicating distinct expression profiles (Fig S4B). Comparison of *nhr-68* mutants and wild-type animals under these basal conditions revealed genes directly regulated by NHR-68 (Table S2). Gene Ontology analysis revealed that genes involved in the innate immune response were highly enriched (Fig S5A and Table S3). Our analysis identified immune genes that overlap with the NHR-68–regulated genes from previously published studies [37, 44], as well as additional immune-related genes uniquely detected in our study (Table S3), despite differences in experimental conditions highlighted in Table S3. Many of these NHR-68–dependent immune genes overlapped with immune-related gene sets previously associated with the PMK-1 and DAF-16 pathways, as well as with the ELT-2 and SKN-1 pathways (Fig S5B and Table S4), consistent with the overlap patterns observed in *P. aeruginosa*–infected samples. These results suggest that NHR-68 represses a subset of immune genes even in the absence of pathogen exposure and functions as a negative regulator of multiple immune signaling pathways.

We hypothesized that if PMK-1 and DAF-16 activation drives the enhanced survival of *nhr-68(gk708)* animals during *P. aeruginosa* infection, then impairing these pathways should reduce the phenotype. Indeed, RNAi knockdown of *pmk-1* or *daf-16* reduced the magnitude of the increased survival in *nhr-68(gk708)* animals infected with *P. aeruginosa* (Figs 3D–3F). These results indicate that the activation of the PMK-1 and DAF-16 immune pathways contributes to the increased pathogen resistance observed in *nhr-68(gk708)* animals.

Having shown that NHR-68 inhibits the PMK-1 and DAF-16 pathways, we next compared NHR-68–dependent gene sets with *P. aeruginosa*-regulated transcriptomes (Table S2 and S3). This analysis revealed overlaps with both infection-induced and infection-repressed gene categories, as well as with immune and metabolic regulatory transcriptional programs, including multiple nuclear hormone receptors. These shared transcriptional features further support a role for NHR-68 in coordinating immune and metabolic gene regulation during pathogen response. Consistent with this transcriptional response, qRT–PCR analysis showed that both *nhr-68* and *nhr-10* transcript levels were modestly increased upon exposure to *P. aeruginosa* in wild-type animals (Fig S6A), indicating that pathogen infection also modulates NHR expression.

To further examine the relationship between NHR-10 and NHR-68 in immune gene regulation, we quantified additional NHR-68–regulated immune genes in *nhr-10(tm4695)* animals (Fig S6B). Analysis of shared target genes revealed that most were upregulated, whereas a subset was unaffected, indicating partial overlap between NHR-10– and NHR-68–mediated immune gene regulation.

### NHR-68 regulates avoidance behavior through polyunsaturated fatty acid metabolism

Pathogen avoidance can be strongly induced by disrupting the defecation motor program, and *aex-5* RNAi animals, which undergo intestinal distension, robustly avoid *P. aeruginosa* [19, 20]. A gas chromatography-mass spectrometry (GC–MS)–based metabolomic profiling we had carried out on these strong-avoiding animals indicated that linoleic acid (LA), an omega-6 polyunsaturated fatty acid, was markedly elevated relative to controls (Fig 4A, Table S5).

**Fig 4.**
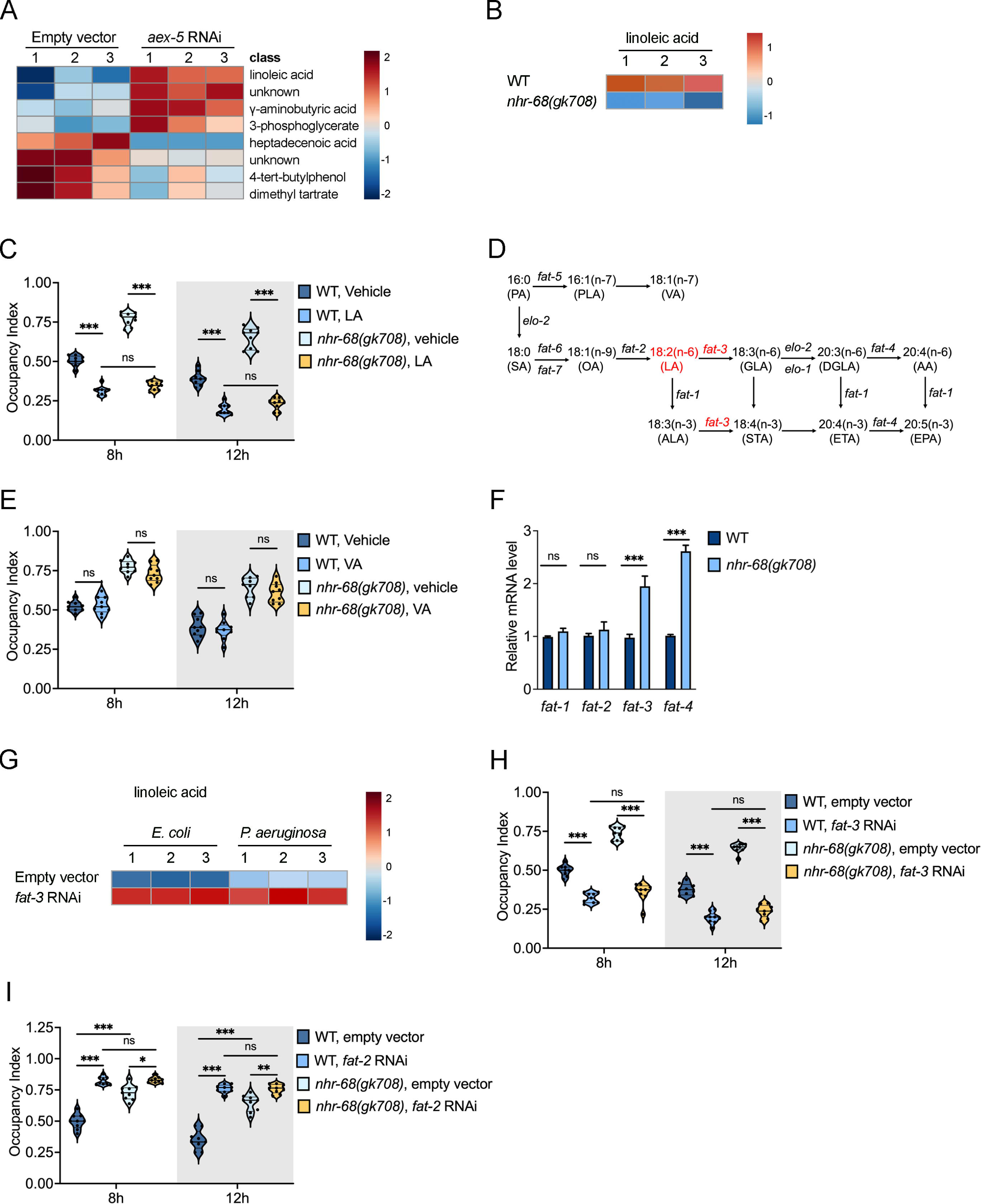
LA accumulation enhances pathogen avoidance. (A) Metabolomic analysis. Animals fed on empty vector or *aex-5* RNAi were harvested for metabolomic analysis using GCOMS. The top-ranked differentially abundant metabolites were identified, and their relative expression was plotted in a heatmap. (B) Relative linoleic acid content measured by LC-HRMS in wild-type and *nhr-68(gk708)* mutant animals is shown as a heatmap; each column represents one of three biological replicates. (C) *P. aeruginosa* occupancy indexes in wild-type and *nhr-68(gk708)* animals treated with vehicle or LA at 8 hr and 12 hr. The occupancy index was calculated as (N _on_ lawn/N _total_). LA, linoleic acid. The bars represent the means, while the error bars indicate the SDs of three independent experiments; ***p<0.001. (D) Schematic of the PUFA synthesis pathway in *C. elegans*, adapted from [29]. The diagram depicts all known steps in PUFA synthesis as well as the elongase and desaturase enzymes involved. Abbreviations: PA, palmitic acid; PLA, palmitoleic acid; VA, vaccenic acid; SA, stearic acid; OA, oleic acid; LA, linoleic acid; ALA, alpha-linolenic acid; GLA, gamma-linolenic acid; STA, stearidonic acid; DGLA, dihommo gamma-linolenic acid; ETA, eicosatrienoic acid; AA, arachidonic acid; EPA, eicosapentaenoic acid. (E) *P. aeruginosa* occupancy indexes in wild-type and *nhr-68(gk708)* animals treated with vehicle or VA at 8 hr and 12 hr. The occupancy index was calculated as (N _on_ lawn/N _total_). LA, linoleic acid. The bars represent the means, while the error bars indicate the SDs of three independent experiments. (F) qRTOPCR analysis of *fat-1*, *fat-2*, *fat-3* and *fat-4* fed on *E. coli*. Data are presented as the means ± SDs from three independent experiments, n = 200 per condition. \*\*\**p* < 0.001, *t* test. (G) GC–MS analysis of linoleic acid abundance in *fat-3* RNAi animals. Animals fed empty vector or *fat-3* RNAi were harvested under *E. coli* or *P. aeruginosa* conditions and subjected to GC–MS–based metabolomic profiling. (H) *P. aeruginosa* occupancy indexes of wild-type and *nhr-68(gk708)* animals fed empty vector control or *fat-3* RNAi at 8 hr and 12 hr. (I) *P. aeruginosa* occupancy indexes of wild-type and *nhr-68(gk708)* animals fed empty vector control or *fat-2* RNAi at 8 hr and 12 hr.

Because NHRs play an important role in regulating lipid metabolism and maintaining fatty acid homeostasis [21, 22, 45], we measured LA levels in *nhr-68(gk708)* animals. Liquid chromatography–high-resolution mass spectrometry (LC-HRMS) analysis showed a significant decrease in LA content in *nhr-68(gk708)* animals (Fig 4B, Table S5). To assess whether elevated LA levels contribute to pathogen avoidance, we supplemented wild-type animals with LA and observed enhanced avoidance behavior toward *P. aeruginosa* (Figs 4C and S7A). We ruled out a direct aversive effect of LA using bacterial choice assays (Fig S7B). To test whether LA is required for NHR-68–mediated avoidance, we supplemented *nhr-68(gk708)* animals with LA. While these *nhr-68(gk708)* animals typically exhibit reduced avoidance behavior, LA treatment fully rescued this defect and further enhanced avoidance behavior. The enhanced avoidance observed in *nhr-68(gk708)* animals was comparable to that of wild-type animals supplemented with LA (Figs 4C and S7A). Importantly, despite its strong effects on pathogen avoidance, LA supplementation did not alter the increased survival phenotype of *nhr-68(gk708)* animals (Figs S8A), indicating that LA-dependent behavioral changes are not sufficient to modify survival outcomes. To examine whether this effect was specific to LA, we tested vaccenic acid (VA; 18:1 n-7), a monounsaturated fatty acid structurally similar to LA but outside the LA metabolic pathway (Fig 4D). VA supplementation did not affect general avoidance behavior or NHR-68–regulated avoidance behavior (Figs 4E and S7C).

To determine if *nhr-68* influences LA metabolism, we measured the expression of genes involved in LA biosynthesis. In *nhr-68(gk708)* animals, we observed elevated expression of *fat-3* and *fat-4*, which are involved in downstream desaturation steps, while *fat-1* and *fat-2*, responsible for LA synthesis (Fig 4D), showed no significant changes (Fig 4F). This expression pattern implies a disruption in LA accumulation in the absence of functional NHR-68.

To test whether LA is controlled by the desaturase pathway downstream of NHR-68, we examined LA in *fat-3* RNAi animals. Based on the PUFA biosynthetic pathway (Fig 4D), *fat-3* knockdown is predicted to cause LA accumulation due to reduced downstream desaturation. GC–MS profiling revealed a significant increase in LA levels upon *fat-3* knockdown (Fig 4G, Table S5), consistent with FAT-3 functioning in LA utilization downstream of NHR-68.

Behaviorally, *fat-3* RNAi, which increases LA levels (Fig 4G), also enhanced pathogen avoidance in wild-type animals (Figs 4H and S7D), linking LA accumulation to avoidance behavior. Importantly, *fat-3* RNAi did not increase the survival of *nhr-68(gk708)* animals (Fig. S8B), further supporting that LA-associated behavioral regulation is separable from survival regulation in this context.

To examine whether FAT-3 also mediates pathogen-associated changes in LA homeostasis, we quantified LA levels by GC-MS in vector control and *fat-3* RNAi animals under both *E. coli* and *P. aeruginosa* conditions. *P. aeruginosa* exposure increased LA abundance in vector control animals, whereas no further increase was observed in *fat-3* RNAi animals, which already displayed elevated basal LA levels (Fig 4G and Table S5). These data are consistent with a role for FAT-3 in regulating LA levels during infection.

Additionally, reduction of LA synthesis by *fat-2* RNAi significantly decreased pathogen avoidance, phenocopying the defect observed in *nhr-68(gk708)* animals (Fig. 4I and S7E). Together, these findings identify LA as a key metabolic component of the NHR-68-regulated fatty-acid pathway that promotes pathogen-avoidance behavior and link intestinal lipid homeostasis to behavioral immune regulation.

### NHR-68 controls pathogen avoidance via the chemosensory neuron AWC

Previous studies have shown that chemosensory neurons AWB, AWC, and ASI are required for *P. aeruginosa*-induced avoidance behavior [20]. Consistent with these findings, genetic ablation of AWC, AWB, or ASI significantly reduced avoidance behavior in the corresponding control animals (Figs 5A–5C and S9A-S9C, empty vector groups). To identify the specific neurons involved in NHR-68-regulated avoidance of *P. aeruginosa*, we used animals with genetic ablation of those individual chemosensory neurons and knocked down *nhr-68* by RNAi in each background. ASE(-) animals were used as a negative control, as ASE neurons are required for *E. faecalis*-induced avoidance but not *P. aeruginosa*-induced avoidance [20]. Among the neuron-ablated strains, only AWC(-) animals failed to exhibit an additional reduction in avoidance behavior upon *nhr-68* knockdown (Figs 5A and S9A). Ablation of AWB, ASI, or ASE neurons did not abolish the effect of *nhr-68* knockdown (Figs 5B–5D, S9B–S9D), indicating that the effect of *nhr-68* RNAi is retained in those backgrounds. These results indicate that AWC neurons are required for the NHR-68-dependent component of behavioral immunity against *P. aeruginosa*.

**Fig 5.**
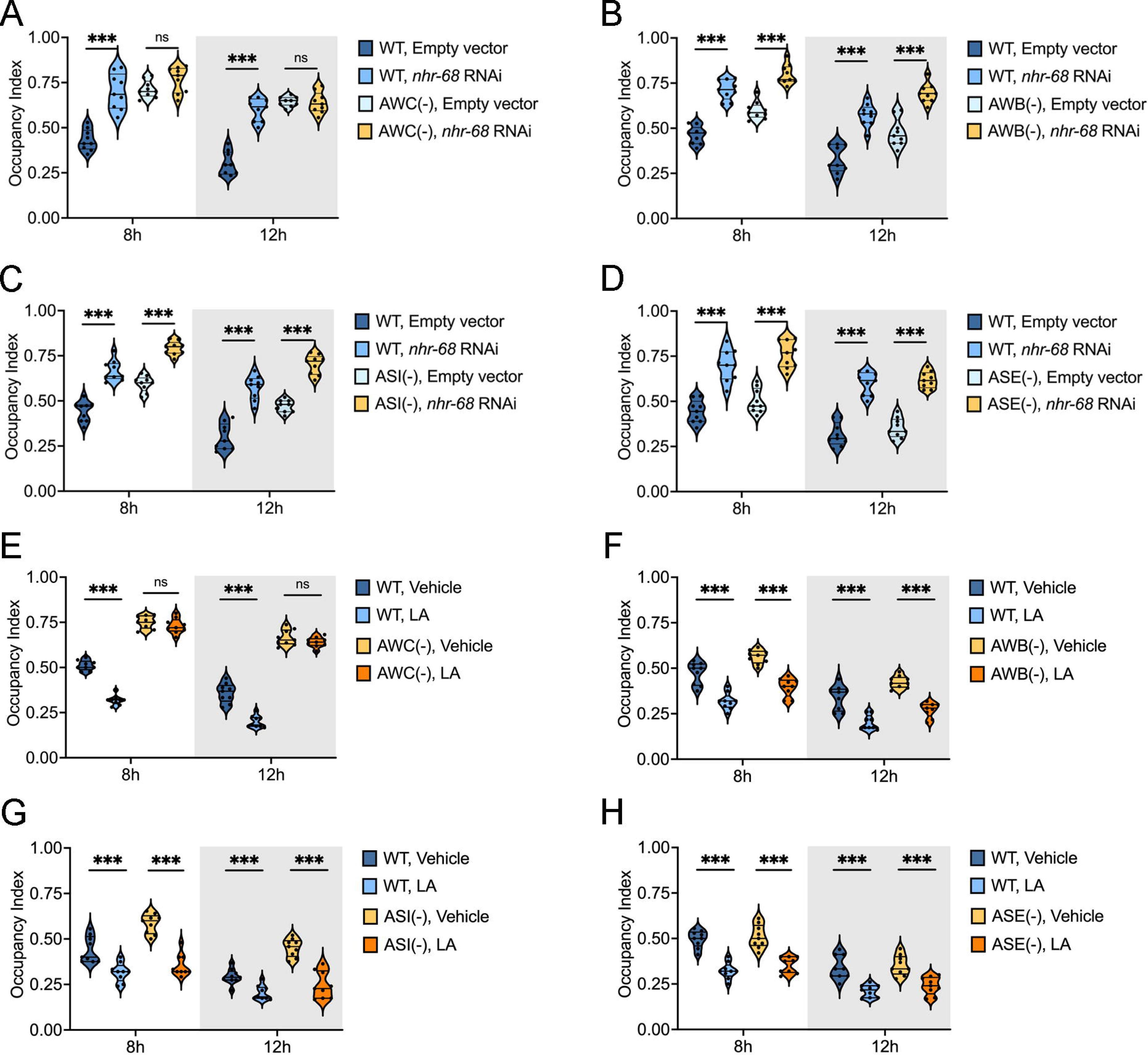
NHR-68 RNAi inhibits pathogen avoidance behavior via the chemosensory neuron AWC. (A) *P. aeruginosa* occupancy indexes of WT and AWC(-) animals fed empty vector control or *nhr-68* RNAi at 8 hr and 12 hr. (B) *P. aeruginosa* occupancy index for WT and AWB(-) animals fed empty vector control or *nhr-68* RNAi at 8 hr and 12 hr. (C) *P. aeruginosa* occupancy indexes of WT and ASI(-) animals fed empty vector control or *nhr-68* RNAi at 8 hr and 12 hr. (D) *P. aeruginosa* occupancy indexes of WT and ASE(-) animals fed empty vector control or *nhr-68* RNAi at 8 hr and 12 hr. (E–H) *P. aeruginosa* occupancy indexes at 8 hr and 12 hr for WT and sensory neuron-ablated animals treated with vehicle or LA. Each panel includes WT animals and one neuron-ablated strain: (E) AWC(-), (F) AWB(-), (G) ASI(-), or (H) ASE(-). LA, linoleic acid. The means and SDs of three independent experiments are shown; the bars, indicated by solid black horizontal lines within the violin plots, represent the means, and the error bars, indicated by thin horizontal lines within the violin plots, represent the SD. *p<0.05, **p<0.01, ***p<0.001, and ns = not significant.

AWC-mediated olfactory adaptation is known to depend on PUFA signaling, and animals with mutations in *fat* genes display defects in AWC function [46]. Since our findings indicate that LA enhances avoidance behavior (Figs 4C and S7A), we next asked whether this effect is also dependent on specific sensory neurons. LA supplementation failed to rescue the avoidance defect in AWC(-) animals, which remained significantly impaired relative to wild-type controls (Figs 5E and S9E), whereas it enhanced avoidance behavior in AWB(-), ASI(-), and ASE(-) animals (Figs 5F–5H and S9F–S9H). Together, these data suggest that AWC neurons are required for both NHR-68-regulated and PUFA-induced avoidance of *P. aeruginosa*, indicating that AWC functions as a specific and consistent component of the behavioral circuit downstream of intestinal metabolic regulation.

Interestingly, reduced avoidance in *nhr-68*-deficient animals did not compromise host defense against *P. aeruginosa*. Instead, *nhr-68(gk708)* animals showed increased survival despite their avoidance defect (Fig 1C). To test whether the neuronal circuitry controlling avoidance also contributes to survival, we examined *nhr-68(gk708)* animals in backgrounds lacking individual chemosensory neurons. Ablation of AWC, AWB, ASI, or ASE neurons did not alter the enhanced survival phenotype of *nhr-68(gk708)* animals on *P. aeruginosa* (Figs S10A-S10D). Under each condition, *nhr-68* RNAi animals remained more resistant than controls. These results indicate that the increased pathogen resistance conferred by loss of *nhr-68* is independent of the chemosensory neurons required for avoidance behavior. Together, these findings suggest that NHR-68 regulates behavioral immunity via AWC neurons, whereas its effect on host survival is mediated through a distinct mechanism.

### The intestinal function of NHR-68 is required for avoidance behavior against *P. aeruginosa*

To identify the tissues in which *nhr-68* functions to regulate avoidance behavior, we performed tissue-specific RNAi knockdown of *nhr-10*, *nhr-68,* or both genes. Intestinal knockdown using the intestinal-specific RNAi strain MGH171 significantly decreased avoidance behavior (Figs 6A and 6B). In contrast, neuron-specific knockdown using strain MAH677, which expresses *rde-1* pan-neuronally and has been previously validated for efficient neuronal RNAi [47, 48], had no effect on avoidance behavior (Fig 6C), indicating that neuronal NHR-68 is not required for behavioral immunity.

**Fig 6.**
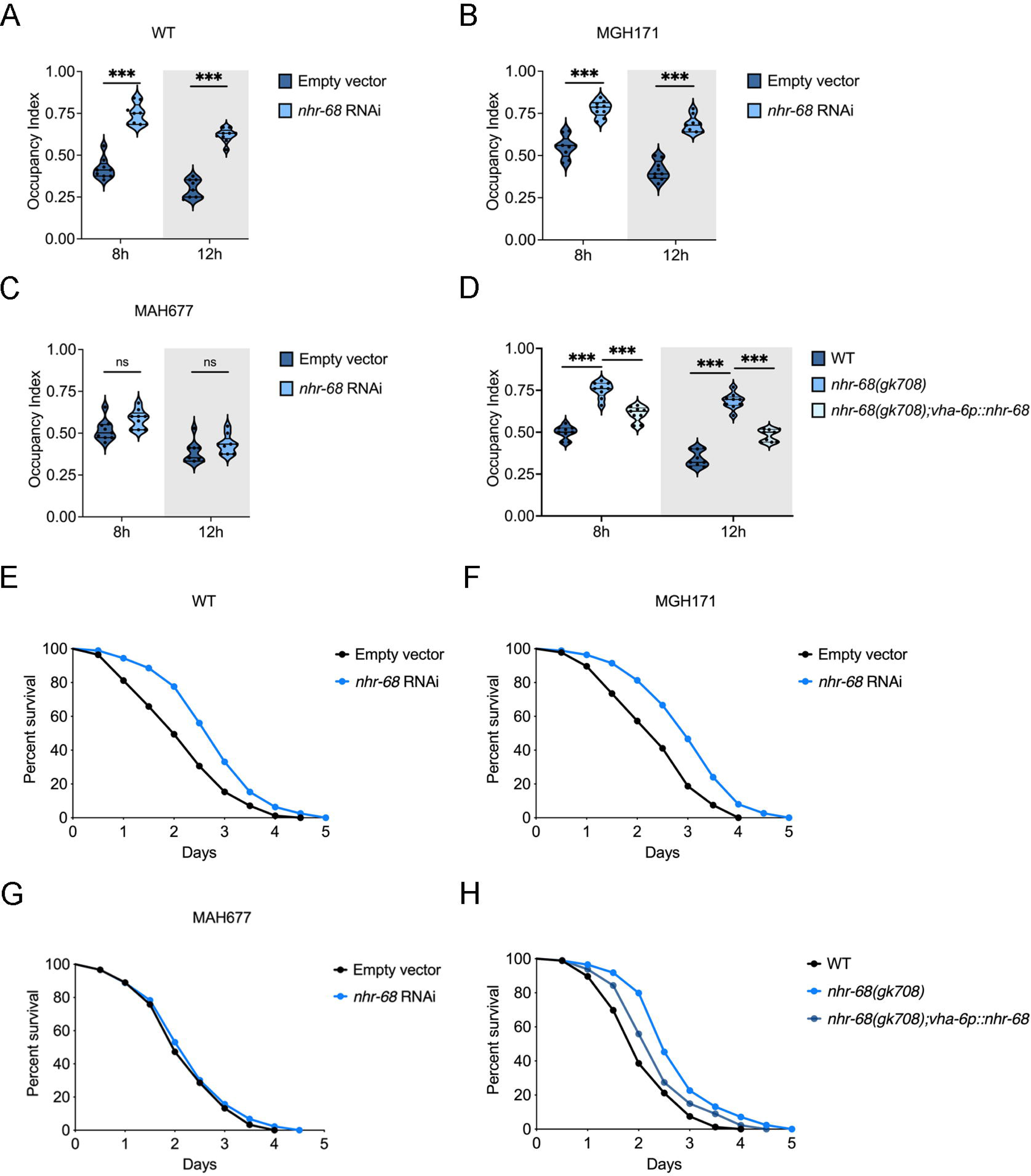
The intestinal function of NHR-68 is required for pathogen avoidance behavior. (A) *P. aeruginosa* occupancy indexes of wild-type animals fed empty vector control or *nhr-68* RNAi at 8 hr and 12 hr. (B) *P. aeruginosa* occupancy indexes of MGH171 animals fed empty vector control or *nhr-68* RNAi at 8 hr and 12 hr. (C) *P. aeruginosa* occupancy indexes of MAH677 animals fed empty vector control or *nhr-68* RNAi at 8 hr and 12 hr. (D) *P. aeruginosa* occupancy indexes for WT, *nhr-68(gk708)* and *nhr-68(gk708);vha-6p::nhr-68* animals at 8 hr and 12 hr after seeding. The occupancy index was calculated as (N _on_ lawn/N _total_). (E) Survival of wild-type animals fed empty vector control or *nhr-68* RNAi on *P. aeruginosa* at 25 °C. Mean survival times were 2.24 and 2.89 days, respectively. Empty vector versus *nhr-68* RNAi, *p* < 0.0001. (F) Survival of MGH171 animals fed empty vector control or *nhr-68* RNAi on *P. aeruginosa* at 25 °C. Mean survival times were 2.23 and 3.02 days, respectively. Empty vector versus *nhr-68* RNAi, *p* < 0.0001. (G) Survival of MAH677 animals fed empty vector control or *nhr-68* RNAi on *P. aeruginosa* at 25 °C. Mean survival times were 2.27 and 2.28 days, respectively. Empty vector versus *nhr-68* RNAi, *p* = ns. (H) Survival of wild-type N2, *nhr-68(gk708)*, or *nhr-68(gk708);vha-6p::nhr-68* animals on *P. aeruginosa* at 25 °C. Mean survival times were 2.17, 2.81, and 2.40 days, respectively. Data are presented from three independent experiments, *n* = 90 per condition. Wild-type versus *nhr-68(gk708)*, p < 0.0001; Wild-type versus *nhr-68(gk708);vha-6p::nhr-68*, *p* = 0.0145; *nhr-68(gk708)* versus *nhr-68(gk708);vha-6p::nhr-68*, *p* = 0.0017.

Since intestinal knockdown of *nhr-68* impaired avoidance behavior, we next tested whether intestine-specific expression of *nhr-68* is sufficient to rescue the reduced avoidance behavior of *nhr-68(gk708)* animals. To drive expression specifically in the intestine, we placed *nhr-68* under the control of the *vha-6* promoter, which is active in intestinal cells [49]. Intestinal expression of *nhr-68* rescued the reduced avoidance behavior of *nhr-68(gk708)* animals (Fig 6D).

Consistent with these findings, intestine-specific knockdown of *nhr-68* also increased survival on *P. aeruginosa*, whereas neuron-specific knockdown had no detectable effect (Figs 6E -6G), indicating that intestinal NHR-68 is required for both behavioral and survival responses during infection. In addition, intestinal expression of *nhr-68* partially rescued the pathogen susceptibility of *nhr-68(gk708)* animals (Fig 6H), further supporting a major role of intestinal NHR-68 in host defense.

Together, these findings indicate NHR-68 functions in the intestine to regulate avoidance behavior against *P. aeruginosa*.

## Discussion

In this study, we identify NHR-68 as a transcriptional regulator that coordinates metabolism and immune responses in *C. elegans* (Fig 7). Our findings show that NHR-68 controls LA metabolism in the intestine and that this regulation influences an AWC-dependent pathogen-avoidance behavior. These results *define* a transcriptional circuit linking intestinal lipid metabolism to the neural control of behavioral immunity, expanding our understanding of how metabolic state influences host defense.

**Fig 7.**
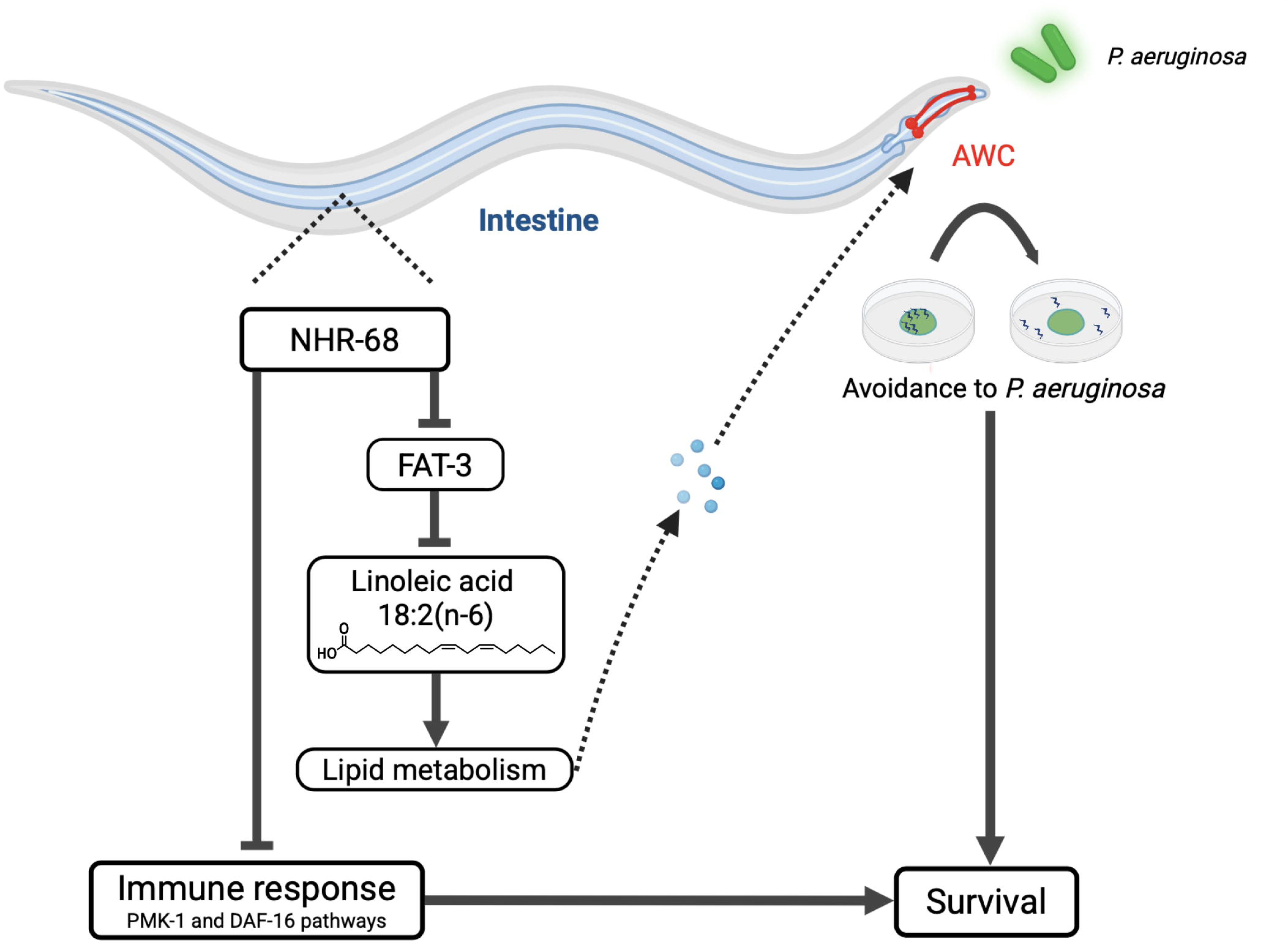
NHR-68 integrates lipid metabolism with behavioral and molecular immunity. NHR-68 acts in the intestine to regulate linoleic acid homeostasis through control of *fat-3* expression. Changes in intestinal lipid metabolism influence an AWC-dependent pathogen-avoidance circuit through currently unidentified intestine-to-neuron signaling mechanisms, enhancing survival during *P. aeruginosa* infection. In parallel, NHR-68 suppresses PMK-1/p38 MAPK and DAF-16/FOXO immune pathways, coordinating behavioral and molecular defenses to optimize host protection.

By regulating PUFA levels, NHR-68 modulates the organism’s ability to mount behavioral responses to pathogens. Fatty acids, particularly PUFAs, are integral to the regulation of various biological processes, including cellular signaling, immune modulation, and membrane dynamics [28, 50]. Our data reveal that alterations in intestinal PUFA metabolism directly affect the ability of *C. elegans* to detect and avoid pathogenic bacteria. This finding is consistent with prior studies showing that metabolic changes can influence behavioral responses to environmental stressors, including infection [51]. Although *fat-3* expression is elevated in *nhr-68(gk708)* animals, it remains unclear whether NHR-68 regulates *fat-3* through direct promoter binding or indirectly through additional transcriptional regulators. Future studies will be required to define the molecular mechanisms underlying this regulation.

We demonstrate that AWC neurons are required for both NHR-68-regulated and LA-enhanced avoidance behavior. However, our data do not identify the molecular signal that links intestinal lipid metabolism to AWC-dependent behavioral responses. While LA supplementation and fatty-acid desaturase perturbations strongly implicate intestinal PUFA homeostasis in this process, future studies will be required to determine whether the relevant signal is LA itself, a lipid-derived metabolite, a neuroendocrine factor, or another intestine-to-neuron signaling mechanism. This connection supports the idea that neural circuits can act as interfaces between physiological states and immune responses. Recent studies have further highlighted the importance of peripheral-to-neuron signaling in coordinating metabolism and behavior. Emerging evidence suggests that metabolic and physiological states in peripheral tissues, including the intestine, can modulate neuronal activity and behavioral outputs through diverse signaling mechanisms [52–54]. In particular, lipid-derived signals and metabolic intermediates have been shown to influence sensory neuron function and behavioral decision-making [55]. These findings provide a conceptual framework for our observations and support a model in which intestinal lipid metabolism, regulated by NHR-68, can shape AWC-dependent pathogen avoidance behavior through periphery-to-neuron communication. PUFAs have been shown to influence chemosensory and olfactory neuron function, including AWC-mediated adaption [46]. Previous work demonstrated that FAT-3 functions cell autonomously in AWC neurons to regulate olfactory adaptation [46], suggesting that PUFA metabolism can directly influence sensory neuron function. Our data further support an additional non-cell-autonomous mechanism in which changes in intestinal lipid metabolic state influence an AWC-dependent behavioral circuit through intestine-to-neuron communication.

In addition to modulating behavior, NHR-68 regulates the activation of the p38/PMK-1 and FOXO/DAF-16 pathways, two conserved regulators of innate immunity. The enhanced expression of immune effectors in *nhr-68* mutants, coupled with the suppression of pathogen resistance by *pmk-1* or *daf-16* RNAi, indicates that NHR-68 functions upstream of these signaling modules. This regulatory role suggests that NHR-68 coordinates metabolic state with immune activation. Beyond PMK-1 and DAF-16, innate immune responses in *C. elegans* are also shaped by additional conserved pathways, including the insulin/IGF-1 signaling (IIS) axis and the SKN-1/Nrf2 oxidative stress response pathway [56, 57], which together integrate metabolic and stress cues to fine-tune host defense. By tempering overactive immune responses, NHR-68 may help balance resource allocation during infection--a principle potentially conserved across species. Such a regulatory balance may prevent the detrimental effects of chronic immune activation, especially in nutrient-limited environments.

In exploring immune gene regulation, we found that NHR-10 and NHR-68 act additively to repress target genes. Our qRT-PCR and reporter analyses show that the double knockdown induces higher *clec-60* expression than either single knockdown, demonstrating that both NHRs independently contribute to immune gene repression.

This additive effect supports their joint contribution to the regulation of molecular and behavioral immune responses. This behavior likely reflects a context-dependent mechanism, where NHR-10 and NHR-68 respond to distinct upstream signals and may target partially overlapping genes with different transcriptional cofactors, highlighting the versatility of nuclear hormone receptor networks in coordinating immune responses.

NHR-10 and NHR-68 have been previously described as a transcriptional feed-forward module in the context of vitamin B12 and propionate metabolism, where NHR-10 acts upstream of NHR-68 to form an AND-logic regulatory circuit required for metabolic gene regulation [37]. In this study, we extend the functional relevance of this conserved module to immune regulation. NHR-10 and NHR-68 jointly regulate pathogen avoidance behavior and immune gene *clec-60* expression. Notably, although NHR-10 functions upstream of NHR-68 at the transcriptional level, their effects on immune outputs are additive, suggesting that feed-forward regulatory wiring can generate graded or combinatorial transcriptional outputs depending on physiological context. This framework provides a mechanism by which conserved nuclear receptor modules integrate metabolic signals and immune regulation, while allowing flexible tuning of downstream functional responses such as behavioral immunity and pathogen resistance.

In addition to NHR-10 and NHR-68, our RNAi screen also identified several other NHRs (e.g., *nhr-77*, *nhr-130*, *nhr-222*, *nhr-162*, *nhr-42*, *nhr-140*, and *nhr-54*) whose knockdown enhanced immune reporter expression. The majority of these receptors remain poorly characterized at the functional level. However, as members of the expanded nuclear hormone receptor family *in C. elegans*, they are likely to participate in broader metabolic and stress-responsive regulatory networks. Their identification in our screen raises the possibility that additional, previously uncharacterized NHRs may contribute to coordinating metabolic state with intestinal immune responses. While genome-wide transcriptomic comparison between *nhr-10* and *nhr-68* mutants remains an interesting direction for future studies, our current genetic and targeted transcriptional analyses indicate that NHR-10 and NHR-68 have partially overlapping roles in coordinating metabolic state and immune regulation.

In wild-type animals, NHR-10 and NHR-68 likely function as metabolic sensors, responding to propionate accumulation and vitamin B12 availability. While their specific ligands remain unidentified, their behavior is consistent with ligand-dependent transcription factors. Upon infection, metabolic stress or altered nutrient absorption could modulate NHR activity, potentially shifting the balance of propionate detoxification and lipid metabolism. Further studies are needed to determine whether infection directly influences NHR-10/NHR-68 activity or ligand availability.

This intersection between lipid metabolism, immune regulation, and behavior is especially notable given the well-established role of PUFAs in inflammation and immunity [58, 59]. Previous studies have also shown that dietary PUFA supplementation can reshape fatty-acid composition and influence host physiology in *C. elegans*, further supporting the idea that changes in PUFA availability can have broad functional consequences [30, 60]. While PUFAs have been studied in the context of cytokine signaling and membrane composition, our findings highlight their importance in shaping behavioral immune responses. That avoidance behavior can be modulated by a defined metabolic product and a transcriptional regulator opens new avenues for dissecting how host physiology integrates metabolic cues to tailor immune and behavioral defenses.

The gut-brain axis has gained increasing attention in recent years as a crucial link between the microbiome, metabolism, and behavior [61–63]. Our results identify NHR-68 as a transcriptional regulator that couples intestinal lipid metabolism to immune and neural outputs in *C. elegans*. By tuning polyunsaturated-fatty-acid balance, NHR-68 shapes pathogen-avoidance behavior through AWC neurons while restraining PMK-1 and DAF-16-dependent immune activation. This coupling provides a mechanistic basis for coordinating behavioral and molecular defenses with the energetic demands of infection. Because nuclear receptors and lipid-derived signals are conserved, similar gut-to-brain mechanisms may integrate metabolic and immune states in other animals.

Together, these findings reveal a general principle in which nuclear receptors align metabolic regulation with adaptive host-defense strategies.

## Materials and Methods

### Strains

The *C. elegans* strains were cultured under standard conditions and fed *E. coli* OP50. All strains were maintained on nematode growth medium (NGM) seeded with *E. coli* (OP50). The *C. elegans* strains used include wild-type N2 Bristol, JIN810 *agIs26* [*clec-60p::gfp* + *myo-2p::mCherry*], RB969 *fat-2(ok873)*, VC1527 *nhr-68(gk708)*, MGH171 *alxIs9* [*vha-6p::sid-1::SL2::GFP*] and MAH677 *sid-1(qt9)* V; *sqIs71* [*rgef-1p*::GFP + *rgef-1p*::sid-1], PY7502 *oyIs85* [*ceh36p::TU#813* + *ceh36p::TU#814* + *srtx1p*::GFP + *unc122p::DsRed*] [AWC(-)], JN1715 *peIs1715* [*str1p::mCasp*-*1*+*unc122p::GFP*] AWB(-), PY7505 *oyIs84* [*gpa-4p*::TU#813+ *gcy-27p*::GFP+*unc-122p*::DsRed] [ASI(-)], PR680 *che-1(p680)* [ASE(-)] were obtained from the *Caenorhabditis* Genetics Center (University of Minnesota, Minneapolis, MN). *nhr-10(tm4695)* was obtained from the National Bioresource Project (NBRP), Japan. Detailed information about the strains used is provided in Table S6.

The following bacterial strains were used: *Escherichia coli* OP50, *E. coli* HT115(DE3), *Pseudomonas aeruginosa* PA14, *Enterococcus faecalis* OG1RF. The *E. coli* OP50, *E. coli* HT115(DE3), and *P. aeruginosa* PA14 cultures were grown at 37°C in Luria–Bertani (LB) broth. The *E. faecalis* cultures were grown at 37°C in brain heart infusion (BHI) broth.

### Feeding RNAi NHR screen

RNA interference (RNAi) was employed to induce loss-of-function RNAi phenotypes by feeding nematodes with *E. coli* HT115(DE3) expressing double-stranded RNA (dsRNA) homologous to the indicated target gene [32, 64]. *E. coli* with the appropriate vectors was grown in LB broth containing ampicillin (100 mg/mL) and tetracycline (12.5 mg/mL) at 37°C overnight and seeded onto NGM plates containing 100 mg/mL ampicillin and 3 mM isopropyl b-D-thiogalactoside (IPTG) (RNAi plates). RNAi-expressing bacteria were allowed to grow overnight at 37°C. Gravid adults were transferred to RNAi-expressing bacterial lawns and allowed to lay eggs for 2 hours. The gravid adults were removed, and the eggs were allowed to develop into young adults for downstream assays. *unc-22* RNAi was included as a positive control to account for RNAi efficiency. All RNAi clones were obtained from the Ahringer RNAi library.

### Fluorescence imaging

Animals were anesthetized in M9 salt solution containing 40 mM sodium azide and mounted onto 2% agar pads. The animals were then visualized using a Leica M165 FC fluorescence stereomicroscope. The GFP signal from eight animals per condition was quantified via Image J software. The fluorescence of an entire animal was calculated with the following equation: corrected whole animal fluorescence = integrated density − (area of selected animal × mean fluorescence of background readings).

### C. elegans killing assays with Pseudomonas aeruginosa

The bacterial lawns used for the *C. elegans* killing assays, performed to assess resistance to *P. aeruginosa*, were prepared by spreading 20 µL of *P. aeruginosa* PA14 culture grown overnight at 37°C on the full surface of modified NGM agar plates (0.35% peptone instead of 0.25% peptone) in 3.5-cm diameter plates. The plates were incubated at 37°C for 16 hours and then cooled to room temperature for at least 1 hour before being seeded with synchronized young adult animals. The killing assays were performed at 25°C, and live animals were transferred daily to fresh plates. Survival was evaluated at the times indicated; animals were considered dead if they failed to respond to touch. Each experiment was performed in triplicate (n = 90 animals).

### *P. aeruginosa* avoidance assay

The bacterial lawns were prepared by inoculating individual bacterial colonies into 3 mL of LB broth as mentioned above and growing them for 24 hours on a shaker at 37 °C. Then, 20 µL of the culture was plated onto the center of a 3.5-cm modified NGM plate (3.5% instead of 2.5% peptone) and incubated at 37 °C for 24 hours. Thirty synchronized young gravid adult hermaphrodites grown on *E. coli* OP50 were transferred outside the bacterial lawns and incubated at 25 °C. Then, the number of animals on and off the lawns were counted at the indicated times for each experiment. The occupancy index was calculated as (N _on_ lawn/N _total_). At least three independent experiments were performed, and three 3.5-cm plates were used per trial in each experiment.

### Reversal assay

Young adult worms (N2, *nhr-68(gk708)*, *nhr-10(tm4695)*, and double mutants) were raised at 20°C. Worms were first transferred to unseeded plates to remove residual bacteria and then placed on unseeded NGM plates. After 1 minute acclimation, reversal events were counted over a 3-minute period. A reversal was defined as at least two consecutive head bends during backward movement.

### Choice Assay with Linoleic Acid Supplementation

A 10 mM linoleic acid (LA) stock was prepared by adding 1 µL of pure linoleic acid to 359 µL of 100% ethanol and vortexing thoroughly to ensure complete mixing. For the choice assay, a 200 µL bacterial suspension was prepared using 20× concentrated overnight *E. coli* or *P. aeruginosa* cultures, fatty-acid-free BSA (5 mg/mL stock), S basal or M9 buffer, and LA or vehicle control. For the LA condition, the mixture consisted of 20× concentrated bacteria (50%), BSA stock (50 mg/mL), buffer (29%), and LA stock (1%, final 100 µM). For the control condition, ethanol was added instead of LA at the same final concentration (1%), with all other components kept identical. Each mixture was vortexed briefly and equilibrated at room temperature before use. For the assay, 30 µL of each condition was spotted onto SK plates in symmetrical positions and allowed to dry prior to behavioral testing. Choice Index = (number of animals on *E. coli* + vehicle − number of animals on *E. coli* + LA) / (number of animals on *E. coli* + vehicle + number of animals on *E. coli* + LA). Choice indices for *P. aeruginosa* were calculated similarly.

### Cloning and generation of transgenic *C. elegans* strains

For *nhr-68* rescue, the recombinant plasmid pPD95.77*_nhr-68p::nhr-68*_SL2::*gfp* was constructed by cloning the *nhr-68* gene and its 670-bp upstream sequence into the *Xba*I and *Sma*I sites of the pPD95.77_SL2::*gfp* vector [65]. For NHR-68 gut-specific rescue, the *nhr-68-*encoding sequence was amplified with the primers *nhr-68*_ATG_*SaL*I F and *nhr-68*_TAA_*Sma*I R. The amplified *nhr-68* DNA was cloned under the *vha-6* promoter in the plasmid pPD95.77_*vha-6p*_SL2 between the *Sal*I and *Sma*I sites to generate the expression clone pPD95.77_*vha-6p*_*nhr-68*_SL2. Transgenic strains were created by injecting 25 ng/mL of the plasmids together with 50 ng/mL of the coinjection marker *unc-122p*::*rfp*.

### C. elegans Killing Assays with Enterococcus faecalis

Twenty microliters of log-phase *E. faecalis* cultures were spread over the entire surface of 35 mm brain-heart infusion (BHI) agar plates, with or without 20 nM VB12. Plates were incubated overnight at 37°C and then cooled to room temperature before seeding with synchronized young adult animals. Killing assays were performed at 25°C, and live animals were transferred daily to fresh plates, with or without 20 nM VB12. Plates containing VB12 were protected from light. Survival was evaluated at the indicated time points; animals were considered dead if they failed to respond to touch. Each experiment was performed in triplicate (n = 90 animals).

### RNA isolation and quantitative reverse transcriptionÕPCR (qRTÕPCR)

Young gravid adult wild-type N2 and *nhr-68(gk708)* animals were either infected for 12 h with *P. aeruginosa* PA14 or fed on *E. coli* OP50 at 25°C. After the exposure period, animals were collected, washed three times with M9 buffer, and immediately frozen in QIAzol reagent (Qiagen, Netherlands). Total RNA was extracted using the RNeasy Plus Universal Kit (Qiagen, Netherlands). A total of 6 µg of total RNA was reverse transcribed with random primers using a high-capacity cDNA reverse transcription kit (Applied Biosystems, Foster City, CA).

Quantitative reverse transcriptionOPCR (qRTOPCR) was conducted using the Applied Biosystems one-step real-time PCR protocol with SYBR Green (Applied Biosystems) on a QuantStudio 5 Real-Time PCR System in 96-well plate format. Reaction mixtures of 25 µl were analyzed as outlined by the manufacturer (Applied Biosystems). The relative fold changes of the transcripts were calculated using the comparative cycle threshold (C_T_) (2^-ΔΔCT^) method and normalized to pan-actin (*act-1, act-3,* and *act-4*). The cycle thresholds of the amplification were determined using QuantStudio 5 software (Applied Biosystems). All samples were run in triplicate. The primer sequences are listed in Table S7.

### RNA sequencing and gene expression analysis

RNA sequencing (RNA-seq) was performed for gene expression analysis and transcriptomic studies. Total RNA was isolated from wild-type N2 and *nhr-68(gk708)* animals infected for 12h with *P. aeruginosa* PA14 in three biological replicates. The sample integrity and purity were checked using an Agilent 2100 system. The RNA was sequenced on a HiSeq X sequencing platform with 150 bp paired-end reads. The RNA sequencing libraries were prepared via the NEBNextâ Ultra RNA Library Prep Kit for Illumina (NEB#E7530L). Library preparation and sequencing were performed at the Novogene Genomic Services & Solutions Company, USA.

The RNA sequence data were analyzed via Lasergene DNA star software. The RNA reads were aligned to the *C. elegans* genome (WS271) via the STAR aligner. Read counts were normalized for sequencing depth and RNA composition across all samples. Differential gene expression analysis was then performed on the normalized samples.

Genes exhibiting at least a twofold change were considered differentially expressed. Gene enrichment of Gene Ontology (GO) biological processes was performed via the Database for Annotation, Visualization, and Integrated Discovery (DAVID) (https://david.ncifcrf.gov/). The overlap of the upregulated genes in previously defined pathways and gene sets, including DAF-16-, PMK-1-, and SKN-1-regulated genes, was calculated. The statistical significance of the overlap between two gene sets was calculated with nemates.org/MA/progs/overlap_stats.html. The representation factor represents the number of overlapping genes divided by the expected number of overlapping genes drawn from 2 independent groups.

### Extraction of metabolites for GCÕMS

*C. elegans* pellets of 2000 animals were lyophilized, and 1.0 mg of biomass was weighed and transferred to a microcentrifuge tube. The metabolites were extracted via the MPLEx protocol [66], in brief, by the addition of 470 µl of a mixture of methanol:water (1.0:1.4), followed by homogenization with a pellet pestle and the addition of 530 µl of ice-cold chloroform. The samples were vortexed for 1 min, placed in an ice block for 5 min, and vortexed again. Centrifugation was performed at 4°C for 10 min at 7,500 × g. The upper and lower layers containing polar and nonpolar metabolites were collected and transferred to glass vials, dried under vacuum, and stored at -20°C. For metabolomic analysis, *aex-5* RNAi animals were grown on *E. coli* HT115 and collected without *P. aeruginosa* exposure.

### Derivatization and GCÕMS acquisition

The vials containing the dried metabolite extracts were stored at -20°C and dried *in vacuo* for an additional 30 min immediately prior to derivatization to ensure the complete removal of residual moisture. The dried extracts were chemically derivatized by adding 20 µL of a 30 mg/ml methoxamine hydrochloride solution in pyridine, followed by vortexing and sonication for 30 s. The samples were then incubated for 90 min in a thermomixer (Eppendorf) at 37°C with shaking at 1000 rpm. After this step was completed, the samples were silylated by adding 80 µL of N-methyl-N-trimethylsilyl-trifluoroacetamide (MSTFA) with 1% trimethylchlorosilane and incubation in a thermomixer at 37°C for 30 min with shaking at 1000 rpm.

The samples were analyzed with an Agilent GC 7890A instrument equipped with an HP-5MS column (30 m × 0.25 mm × 0.25 μm; Agilent Technologies, Santa Clara, CA) coupled with a single quadrupole MSD 5975C (Agilent Technologies). The helium gas flow rate was determined by the Agilent Retention Time Locking (RTL) function on the basis of analysis of deuterated myristic acid (Agilent Technologies, Santa Clara, CA).

One microliter of the derivatized sample was injected into a splitless port at a constant temperature of 250°C. The GC temperature gradient started at 60°C and was held at that temperature for 1 minute after injection, followed by an increase to 325°C at a rate of 10°C/minute and a 10-minute hold at this temperature. The transfer line and quadrupole temperatures were 280°C and 150°C, respectively. The electron impact was set to 70 eV, and mass spectra were collected at a rate of 2.6 Hz over a scanning range of m/z 50–600. A fatty acid methyl ester (FAME) mixture (C8-28) (SigmaOAldrich) was analyzed with each batch as a standard for retention time calibration.

### GCÕMS data analysis

Raw GCOMS data files were processed via Metabolite Detector software [67]. The retention indices (RIs) of the detected metabolites were calculated on the basis of the analysis of the FAME standards, followed by chromatographic alignment across all analyses after deconvolution. Metabolites were identified by matching experimental spectra to a PNNL augmented version of the Agilent GCOMS Metabolomics Library, which contains mass spectra and validated RIs for over 1200 metabolites. All metabolite identifications and quantification ions were manually confirmed to reduce deconvolution errors during automated data processing and to eliminate false identifications. The NIST20 and Wiley11 spectral libraries were also used to cross-validate the spectral matching scores obtained using the PNNL augmented library. The unknown peaks were subjected to further matching to the NIST20 and Wiley11 GCOMS libraries and reported as tentative identifications (denoted by an asterisk) if the MS score was high. The reported metabolite peak areas were normalized to the biomass used for extraction. A heatmap was generated using MetaboAnalyst 5.0 (https://www.metaboanalyst.ca/MetaboAnalyst/).

### Analysis of Fatty Acids by LC-HRMS

Worm samples were homogenized in 300 µL of PBS. From each homogenate, 250 µL was mixed with 10 µL of internal standard (oleic acid-d17, 10 µg/mL) and saponified, followed by extraction with 4 mL of hexane. Extracts were vortexed for 10 min and centrifuged at 4,000 g for 10 min at 4°C. Supernatants were transferred to clean glass tubes and evaporated to dryness under nitrogen. Dried samples were reconstituted in 150 µL of isopropanol, and 5 µL was injected into a Thermo Vanquish LC system equipped with a Kinetex C18 column (2.1 × 100 mm, 2.6 µm). Mobile phase A was 0.1% formic acid in water, and mobile phase B was 80:20 acetonitrile:isopropanol. The flow rate was 300 µL/min at 35°C, and fatty acids were separated over a 25-min gradient.

Data were acquired on a Thermo Orbitrap Lumos mass spectrometer in negative ESI mode at a resolution of 120,000 (scan range 100–500 m/z) and processed using Skyline-daily software.

### Fatty acid treatment

Synchronized young adult animals were collected from NGM plates, washed twice with S basal buffer, and transferred to 6 cm dishes containing 5.8 mL of S basal supplemented with 0.02% NP-40 and 180 µL of an overnight *E. coli* OP50 culture. Linoleic acid (LA) or vaccenic acid (VA) was added to a final concentration of 100 µM from 100 mM ethanol stocks pre-mixed with 0.1% fatty acid–free bovine serum albumin (BSA) to enhance solubility and bioavailability [68]. Cultures were gently shaken at 60 rpm for 4 h at room temperature and protected from light. After incubation, animals were washed three times with S basal to remove residual fatty acids before subsequent behavioral assays. Control treatments contained equivalent amounts of ethanol and 0.1% BSA.

### Quantification and Statistical Analysis

Statistical analysis was performed with GraphPad Prism 8 version 8.1.2 (GraphPad). All error bars represent the standard deviation (SD). Two-sample *t* tests were used for comparisons between two groups, while one-way ANOVA followed by Tukey’s multiple-comparison test was applied for comparisons among three or more groups. *P* values < 0.05 were considered statistically significant. In the figures, asterisks (*) denote statistical significance as follows: ns, not significant; *, *p* < 0.05; **, *p* < 0.01; ***, *p* < 0.001, compared with the appropriate controls. The KaplanOMeier method was used to calculate the survival fractions, and the statistical significance of differences between survival curves was determined via the log-rank test. All experiments were performed at least three times.

## Supporting Information

### Supporting Figures

Fig S1. Quantification of fluorescence signals.

Fig S2. *P. aeruginosa* occupancy of wild-type and mutant animals.

Fig S3. Vitamin B12 addition does not change NHR-68—inhibited innate immune response.

Fig S4. Principal Component Analysis (PCA) for gene expression profiles.

Fig S5. NHR-68 inhibits immune pathways.

Fig S6 NHR-10 influences a subset of NHR-68–regulated immune genes and both respond to pathogen exposure.

Fig S7. *P. aeruginosa* occupancy of animals treated with LA.

Fig S8. Survival of wild-type and *nhr-68(gk708)* animals on *P. aeruginosa* with LA accumulation.

Fig S9. NHR-68-controlled pathogen avoidance behavior requires the chemosensory neuron AWC.

Fig. S10. Survival of wild-type and *nhr-68* RNAi animals on *P. aeruginosa* in neuron-ablated backgrounds.

### Supporting Tables

Table S1. Fluorescence Quantification of Screened NHR Genes

Table S2. Upregulated and downregulated genes more than 2-fold (p < 0.05) in non-infected and *P. aeruginosa*-infected *nhr-68(gk708)* versus WT animals

Table S3. GO enrichment analysis

Table S4. Representation factors for upregulated genes in *P. aeruginosa*-infected *nhr-68(gk708)* animals

Table S5. Metabolomic profiling and lipid quantification in NHR-68 pathway

Table S6. Strains

Table S7. Primers

## Acknowledgements

We thank current and former Aballay Lab members for insightful discussions. We thank Meagan Burnet and Nathalie Munoz (Biological Sciences Division, Pacific Northwest National Laboratory, Richland, WA 99352, USA) for their contributions to GC–MS analyses, including quality control and sample processing. Most strains used in this study were obtained from the *Caenorhabditis* Genetics Center (CGC), which is funded by the NIH Office of Research Infrastructure Programs (P40 OD010440) and the National BioResource Project (NBRP), Japan.

