## Supporting Figures for "Intestinal lipid metabolism controls immune response through NHR-68 and gut–brain signaling in *C. elegans*"

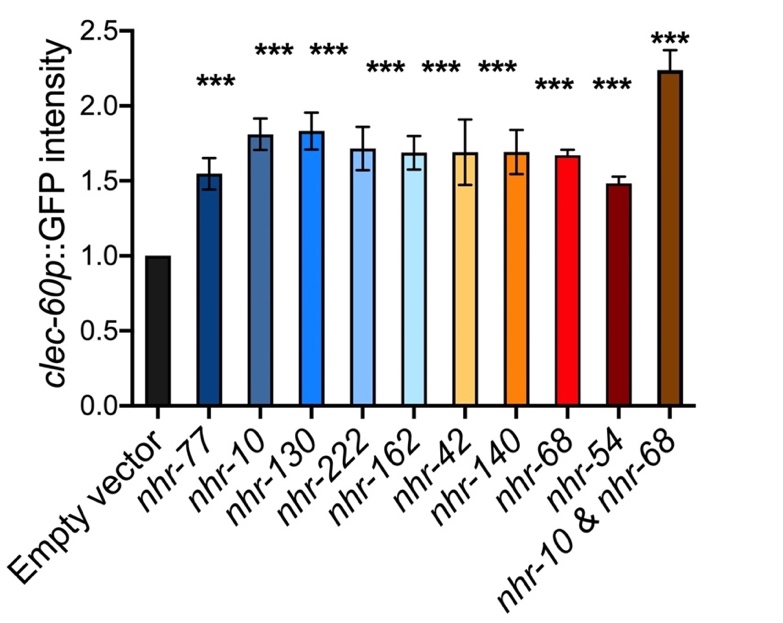

**Fig S1. Quantification of ﬂuorescence signals.** Quantification of ﬂuorescence signals in Fig 1A. n=8. The “n” represents the number of animals in each experiment. *N* = 3 biological replicates.

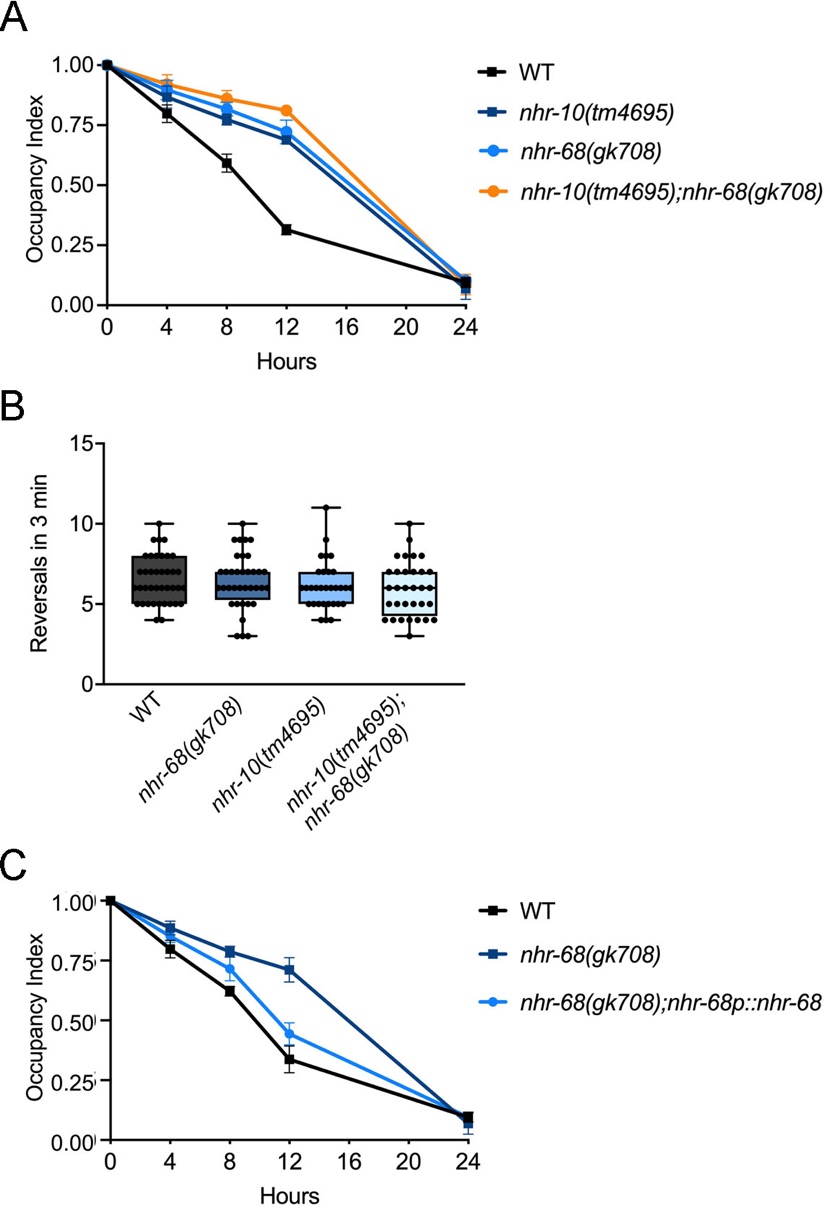

**Fig S2. *P. aeruginosa* occupancy of wild-type and mutant animals.** (A) Time course of the percent occupancy on *P. aeruginosa* of WT, *nhr-10(tm4695)*, *nhr-68(gk708),* and *nhr-10(tm4695)*;*nhr-68(gk708)* animals grown on *P. aeruginosa* partial lawns. WT versus *nhr-10(tm4695)*, *p* < 0.001; WT versus *nhr-68(gk708)*, *p* < 0.001; WT versus *nhr-10(tm4695)*;*nhr-68(gk708)*, *p* < 0.0001, *t* test. (B) Locomotion analysis of *nhr-10(tm4695)* and *nhr-68(gk708)* mutants. Reversal frequency was measured in young adult worms and used as an indicator of locomotor ability. (C) Time course of the percent occupancy on *P. aeruginosa* of WT, *nhr-68(gk708)*, and *nhr-68(gk708);nhr-68p::nhr-68* animals grown on *P. aeruginosa* partial lawns. WT versus *nhr-68(gk708)*, *p* < 0.001; *nhr-68(gk708)* versus *nhr-68(gk708);nhr-68p::nhr-68*, *p* < 0.01, *t* test.

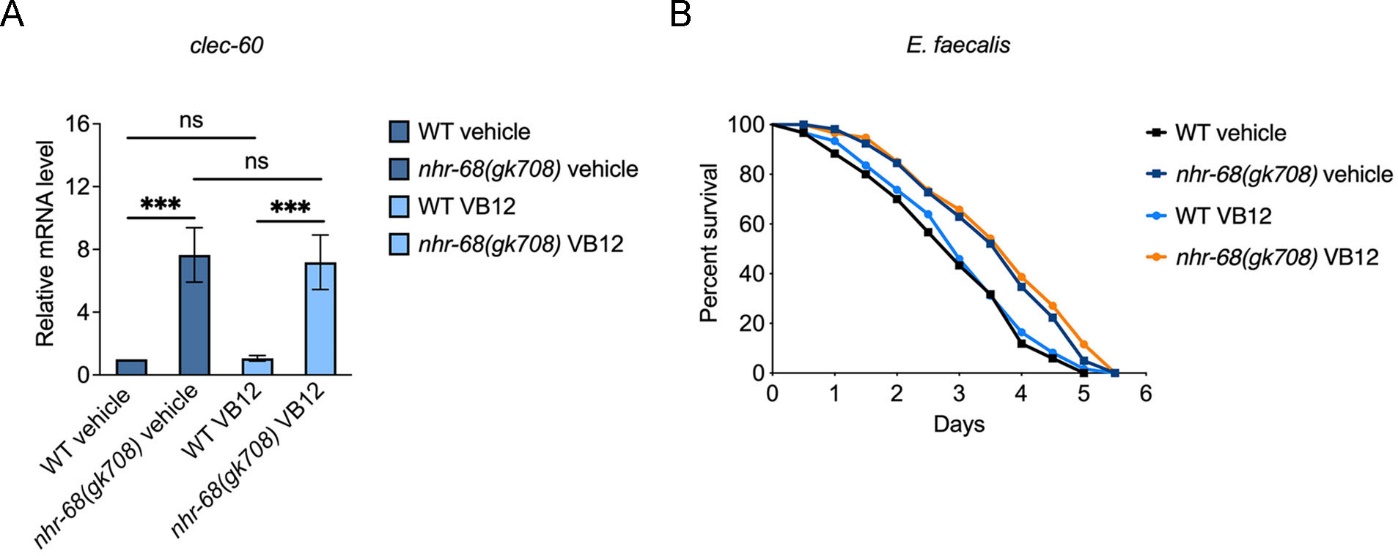

**Fig S3. Vitamin B12 addition does not change NHR-68—inhibited innate immune response.** (A) qRT‒PCR analysis of the immune gene *clec-60* in wild-type and *nhr-68(gk708)* animals fed *E. coli* with or without 20 nM vitamin B12. Data are presented as the means ± SDs from three independent experiments, n = 200 per condition. ****p* < 0.001, one-way ANOVA with Tukey’s multiple-comparison test. (B) Survival of wild-type and *nhr-68(gk708)* animals grown on *E. faecalis* with or without 20 nM vitamin B12 at 25 ˚C. Mean survival times were 2.97, 3.01, 3.26, and 3.28 days, respectively. Data are presented as means from three independent experiments, n = 90 per condition. WT vehicle versus WT VB12, *p* = ns; WT vehicle versus *nhr-68(gk708)* vehicle, *p* = 0.026; WT VB12 versus *nhr-68(gk708)* VB12, *p* =0.0020; *nhr-68(gk708)* vehicle versus *nhr-68(gk708)* VB12, *p* =0.4739. The Kaplan-Meier method was used to calculate the survival fractions, and statistical significance between survival curves was determined using the log-rank test.

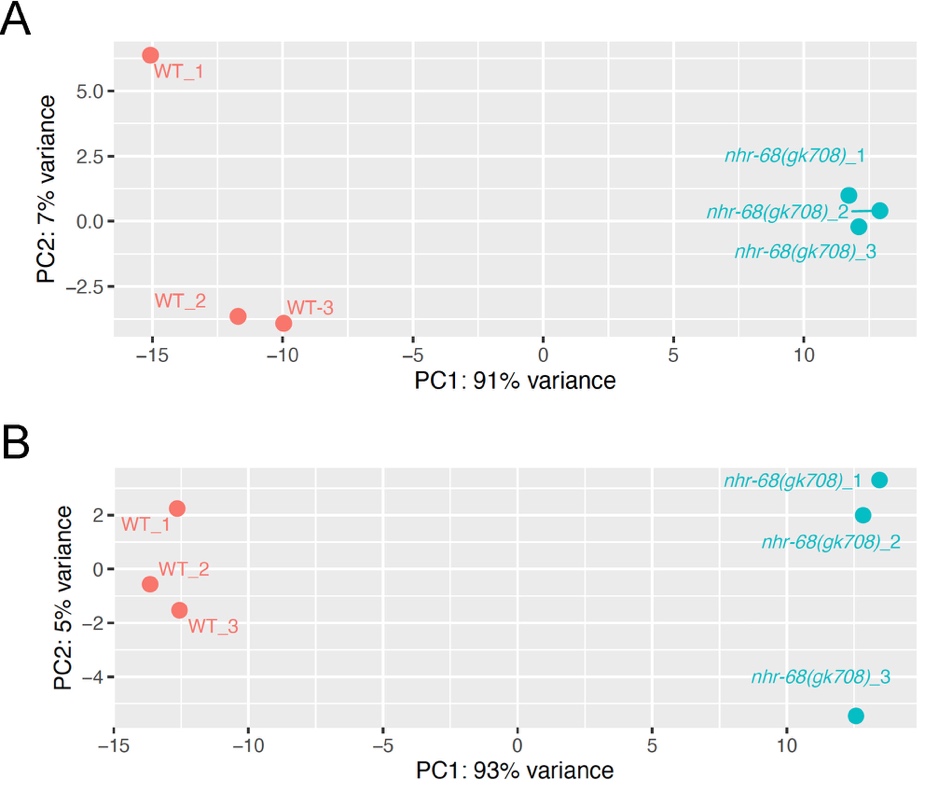

**Fig S4. Principal Component Analysis (PCA) for gene expression profiles.** (A) PCA plot showing transcriptomic profiles of wild-type and *nhr-68(gk708)* animals after infection with *P. aeruginosa*. (B) PCA plot showing transcriptomic profiles of wild-type and *nhr-68(gk708)* animals fed on *E. coli*. Red and green dots indicate wild-type and *nhr-68(gk708)* animals, respectively

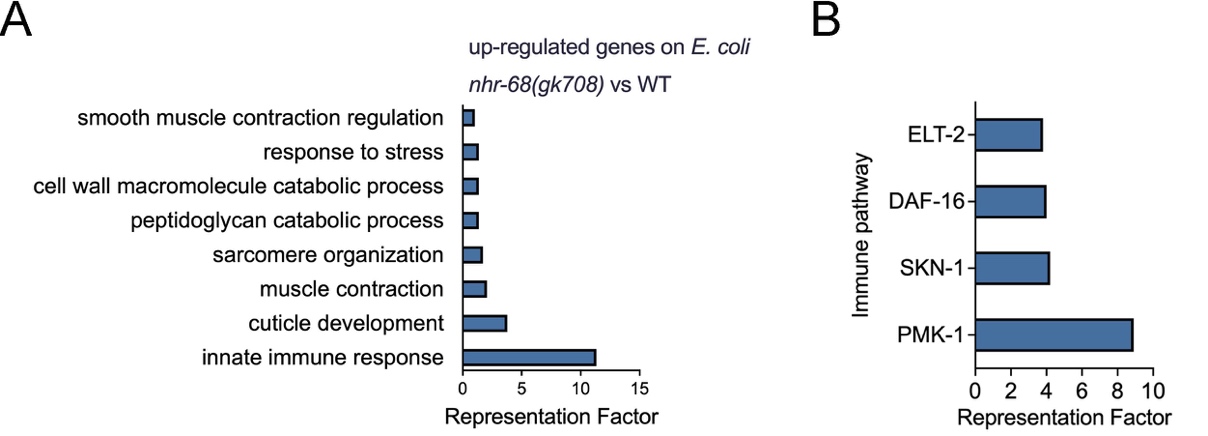

**Fig S5. NHR-68 inhibits immune pathways.** (A) Gene Ontology analysis of genes upregulated in *nhr-68(gk708)* vs. wild-type animals fed on *E. coli* OP50. Only terms significantly enriched in *nhr-68(gk708)* (q < 0.1) are shown. (B) Representation factors of upregulated genes controlled by the PMK-1, SKN-1, DAF-16, and ELT-2 pathways. Representation factors greater than 1 indicate significant overlap with the corresponding pathway gene sets.

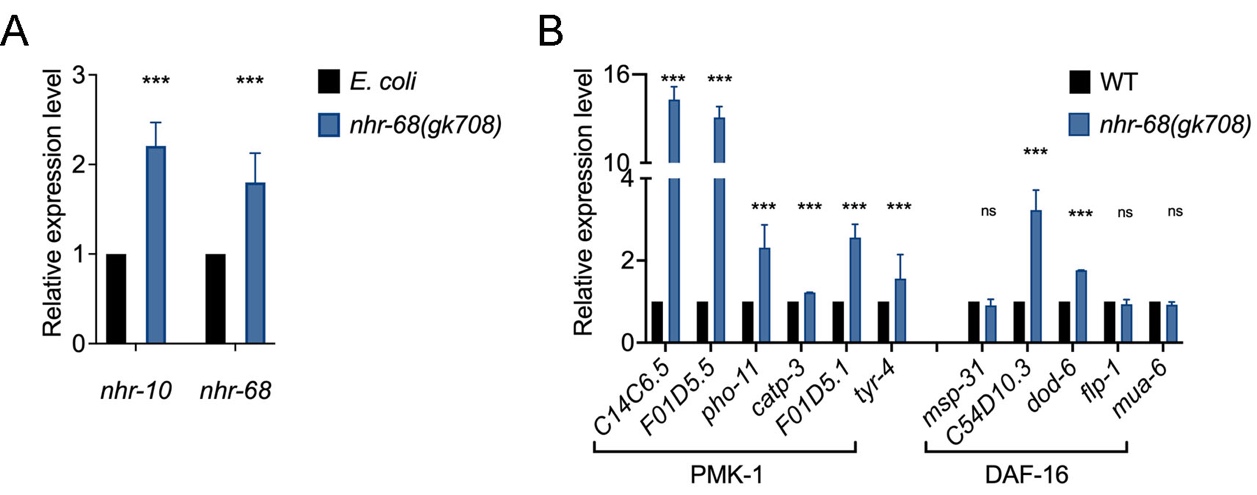

**Fig S6 NHR-10 influences a subset of NHR-68–regulated immune genes and both respond to pathogen exposure.** (A) qRT–PCR analysis of *nhr-10* and *nhr-68* transcript levels in wild-type animals exposed to *P. aeruginosa* relative to *E. coli*-fed controls. (B) qRT–PCR analysis of NHR-68–regulated immune genes shown in Fig. 3C in *nhr-10(tm4695)* animals. *p < 0.05, **p < 0.01, ***p < 0.001, and ns = not significant.

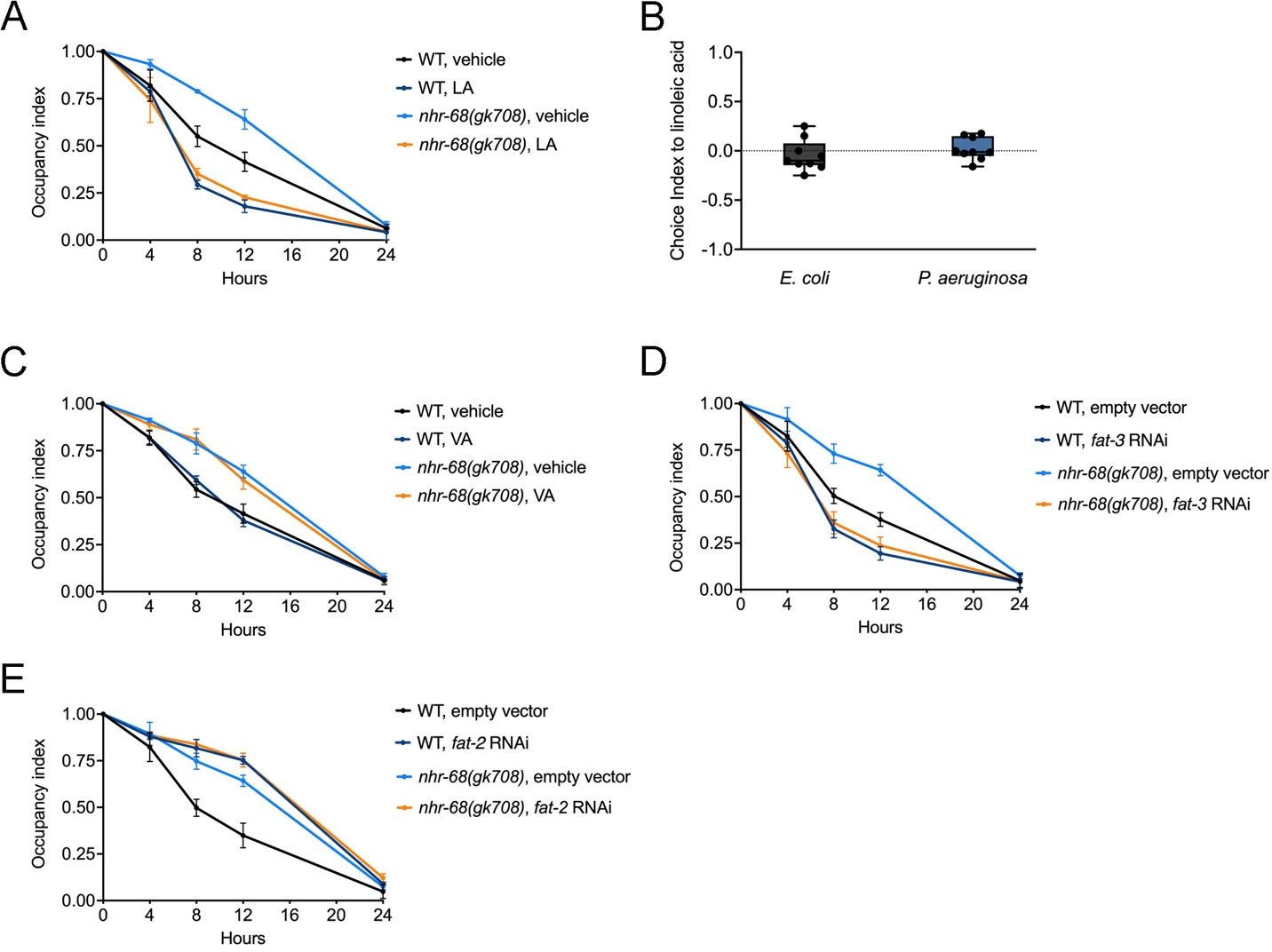

**Fig S7. *P. aeruginosa* occupancy of animals treated with LA.** (A) Time course of the percent occupancy on *P. aeruginosa* of wild-type and *nhr-68(gk708)* animals treated with vehicle or LA. (B) Linoleic acid does not function as a direct aversive cue in bacterial choice assays. Two-choice assays were performed using *E. coli* or *P. aeruginosa* lawns supplemented with linoleic acid (LA) or vehicle. Choice Index (CI) was calculated as (N_vehicle − N_LA)/(N_vehicle + N_LA). No significant preference or avoidance was observed under either condition, indicating that LA does not act as an innate repellent in *C. elegans*. Data are shown as mean ± SD. (C) Time course of the percent occupancy on *P. aeruginosa* of wild-type and *nhr-68(gk708)* animals treated with vehicle or VA. (D) Time course of the percent occupancy on *P. aeruginosa* of wild-type and *nhr-68(gk708)* animals fed empty vector control or *fat-3* RNAi. The occupancy index was calculated as (N _on_ lawn/N _total_). (E) Time course of the percent occupancy on *P. aeruginosa* of wild-type and *nhr-68(gk708)* animals fed empty vector control or *fat-2* RNAi.

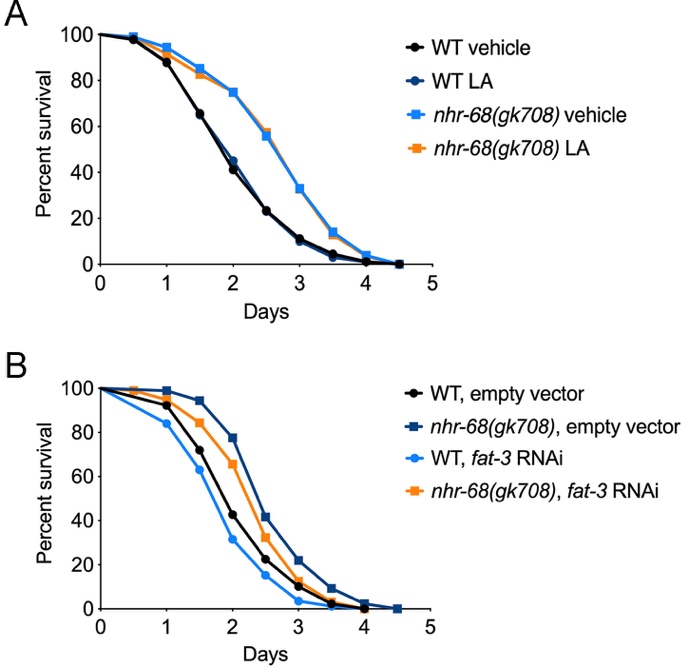

**Fig S8. Survival of wild-type and *nhr-68(gk708)* animals on *P. aeruginosa* with LA accumulation.** (A) Survival of wild-type N2 and *nhr-68(gk708)* animals on *P. aeruginosa* PA14 at 25 °C after linoleic acid treatment. WT vehicle versus WT LA, *p* = ns; *nhr-68(gk708)* vehicle versus *nhr-68(gk708)* LA, *p* = ns. Mean survival times were 2.16, 2.15, 2.56 and 2.55 days, respectively. (B) Survival of wild-type N2 and *nhr-68(gk708)* animals on *P. aeruginosa* PA14 at 25 °C after RNAi treatment with empty vector or *fat-3* RNAi. Mean survival times were 2.07, 2.63, 1.91 and 2.50 days, respectively. WT empty vector versus WT *fat-3* RNAi, *p* = 0.05; *nhr-68(gk708)* empty vector versus *nhr-68(gk708)* *fat-3* RNAi, *p* = 0.0263. Data are presented as means from three independent experiments, n = 90 per condition.

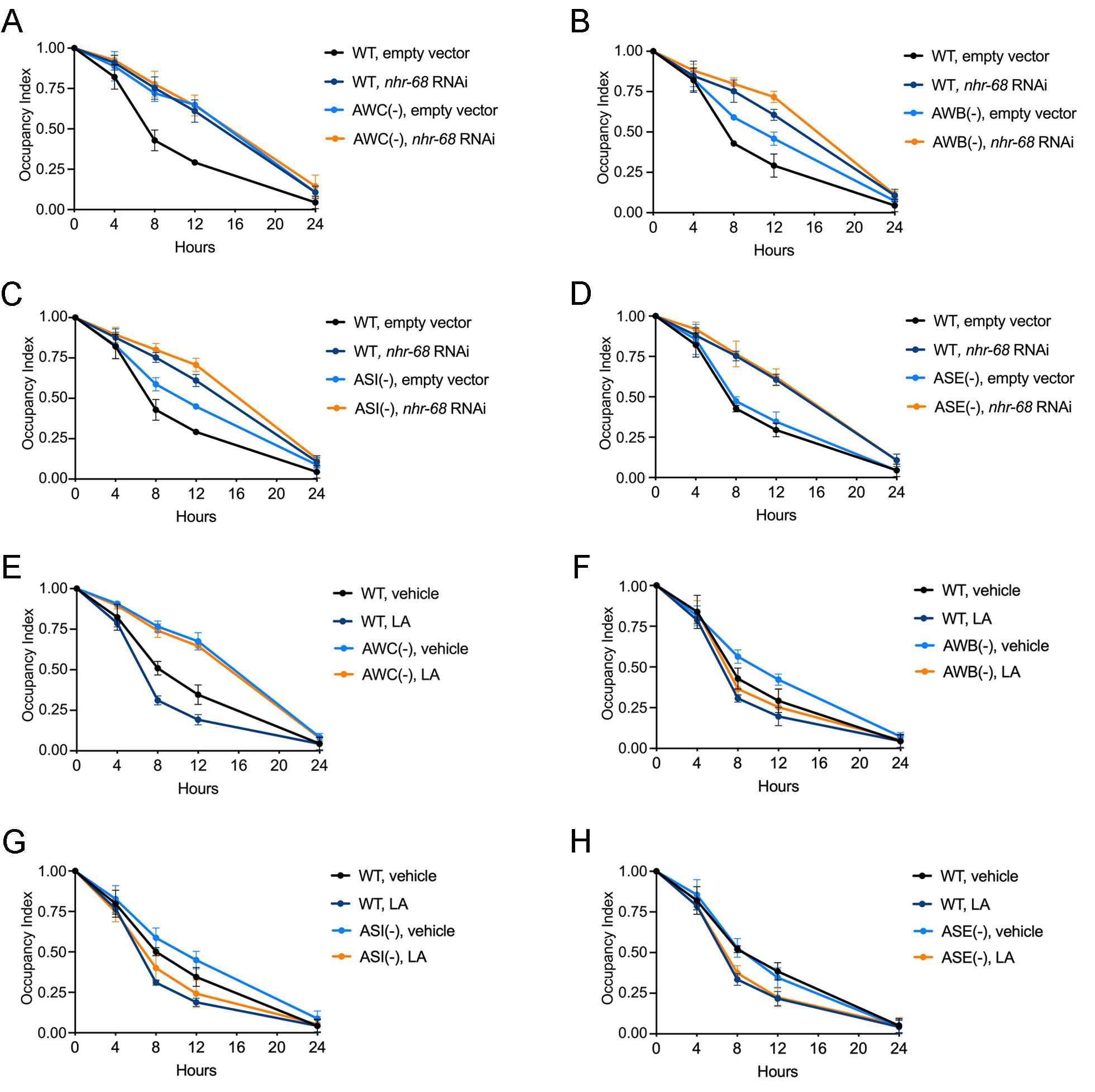

**Fig S9. NHR-68-controlled pathogen avoidance behavior requires the chemosensory neuron AWC.** (A) Time course of the percent occupancy on *P. aeruginosa* for WT and AWC(-) animals fed empty vector control or *nhr-68* RNAi. (B) Time course of the percent occupancy on *P. aeruginosa* for WT and AWB(-) animals fed empty vector control or *nhr-68* RNAi. (C) Time course of the percent occupancy on *P. aeruginosa* for WT and ASI(-) animals fed empty vector control or *nhr-68* RNAi. (D) Time course of the percent occupancy on *P. aeruginosa* for WT and ASE(-) animals fed empty vector control or *nhr-68* RNAi. (E–H) Time course of the percent occupancy on *P. aeruginosa* partial lawns for WT and sensory neuron-ablated animals treated with vehicle or LA. Each panel includes WT animals and one neuron-ablated strain: (E) AWC(-), (F) AWB(-), (G) ASI(-), or (H) ASE(-). LA, linoleic acid.

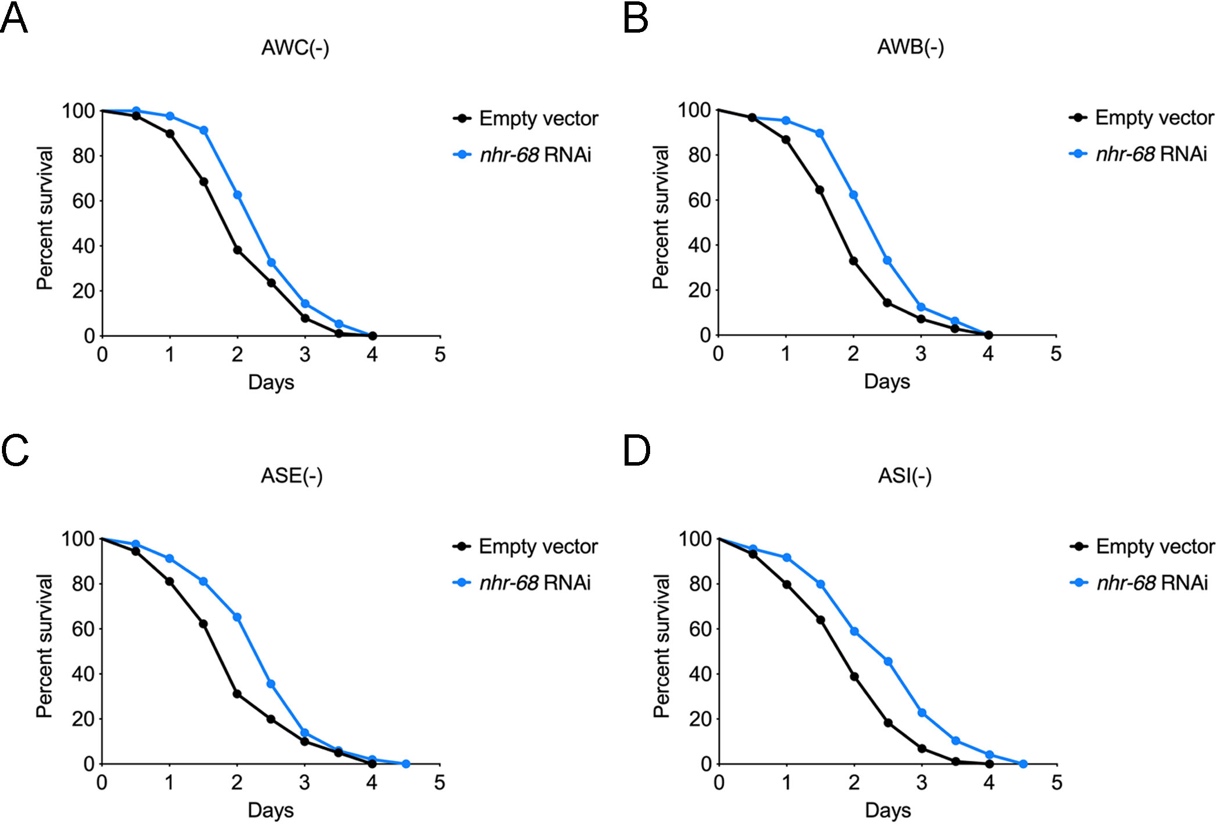

**Fig. S10. Survival of wild-type and *nhr-68* RNAi animals on *P. aeruginosa* in neuron-ablated backgrounds.** (A–D) Survival of wild-type N2 and *nhr-68* RNAi animals on *P. aeruginosa* PA14 at 25 °C in AWC(-), AWB(-), ASE(-), or ASI(-) neuronal ablation backgrounds. Data are presented as means from three independent experiments, n = 90 per condition.
